# Endogenous SNAP-tagging of Munc13-1 for monitoring synapse nanoarchitecture

**DOI:** 10.1101/2024.10.08.617143

**Authors:** Maria Kowald, Sylvestre P. J. T. Bachollet, Fritz Benseler, Maria Steinecker, Moritz Boll, Sofia Kaushik, Tolga Soykan, Siqi Sun, Ramona Birke, Dragana Ilic, Nils Brose, Hanna Hörnberg, Martin Lehmann, Silvio O. Rizzoli, Johannes Broichhagen, Noa Lipstein

**Author notes:** Corresponding Authors **Johannes Broichhagen −** Leibniz-Forschungsinstitut für Molekulare Pharmakologie, Berlin 13125, Germany, **Noa Lipstein –** Max Planck Institute for Multidisciplinary Sciences, Göttingen 37077, Germany, Leibniz-Forschungsinstitut für Molekulare Pharmakologie, Berlin 13125, Germany.

## Abstract

Synaptic function is governed by highly regulated protein machineries, whose abundance and spatial localization change continually. Studies to determine the dynamic changes in synaptic proteins nanoarchitecture typically rely on immunolabeling, or on the expression of fluorescent proteins. The former employs chemical fluorophores and signal amplification, but requires fixation. The latter enables monitoring of proteins in live microscopy, but uses suboptimal fluorophores. Self-labeling tags have been introduced to combine the advantages of these two approaches, and here we introduce a knock-in mouse line where the essential presynaptic protein Munc13-1 is endogenously fused to the self-labeling SNAP tag. We demonstrate efficient Munc13-1-SNAP labeling in fixed neurons and brain sections by various SNAP dyes, as well as by a novel far-red and cell impermeable compound, SBG-SiR-d12. We introduce and characterize SBG-SiR-d12 here as a highly-efficient dye for SNAP-tag labeling of extracellular epitopes, and of intracellular proteins in fixed and permeabilized tissue. Finally, we show that Munc13-1-SNAP can be labeled in living neurons and monitored through live-cell imaging using confocal- and super resolution microscopy. We conclude that the Unc13a^SNAP^ mouse line is a useful tool for the analysis of presynaptic nanoarchitectural dynamics, with a potential for wide adoption.

## INTRODUCTION

Dynamic changes in protein copy numbers, complex composition, and nanoscale organization, often follow alterations in cellular activity levels. To reliably monitor proteins in time and space, efficient labeling strategies have been developed. A key development has been the introduction of fluorescent protein tags (e.g. green fluorescent protein, GFP^1, 2^). However, and despite continuous improvements, the photophysical properties of fluorescent proteins often fall short compared to chemical dyes in terms of brightness and stability^3, 4^. To address this limitation, self-labeling tags were created - engineered protein tags that can be genetically encoded and covalently bind bioorthogonal synthetic probes. Belonging to this group, the SNAP tag^5^ is an engineered *O*^6^-alkylguanine-DNA alkyltransferase that binds *O*^6^-benzylguanine (BG) derivatives in a covalent, non-reversible manner^6–8^. As such, proteins fused to a SNAP tag can be visualized in live cells or in fixed tissue by the addition of a BG-fluorophore conjugate. This mode of protein labeling is advantageous as an alternative to genetic conjugation with fluorescent protein tags, because it enables the flexible attachment of bright and stable dyes to the protein of interest, in the living cell, and at a time of choice. The size of the SNAP tag (∼19 kDa) brings the fluorophore in proximity to the protein of interest^9, 10^, which may be advantageous in super-resolution microscopy^11^. Moreover, flexible use of chemical dyes enables pulse-chase labeling^12–15^ or signal multiplexing^16^. The toolkit for SNAP tag labeling is rapidly expanding: BG derivatives carrying an array of fluorophores are available, and several chemical modifications of the BG-dye conjugates have been introduced to modify their properties, for example to make the conjugate membrane impermeable for extracellular labeling^13^ or to boost labeling kinetics^17^.

Here, we opted to leverage the advantages of self-labeling tags and develop novel tools to monitor synapses. Munc13-1 (protein unc-13 homolog A, encoded by the *Unc13a* gene) is a presynaptic protein with a central function in the preparation of synaptic vesicles (SVs) for fast exocytosis^18, 19^. Munc13-1 is expressed in the majority of neuronal subtypes in the central and peripheral nervous system, as well as in some neurosecretory cell types including chromaffin cells or insulin-releasing beta-pancreatic cells^20, 21^. In neurons, it resides in the active zone, a protein-dense compartment at the presynaptic membrane where SVs undergo fusion, and is absolutely essential for synaptic transmission. At the molecular level, Munc13-1 catalyzes the formation of SNARE complexes, that link the SV membrane with the presynaptic plasma membrane, thus making SVs fusion-competent^22, 23^. Munc13-1 function sets multiple synapse properties, including synaptic strength, the release probability of SVs, the rate of SV replenishment after depletion, and synaptic plasticity^19, 23–32^. In humans, genetic variations in the *UNC13A* gene are associated with a neurodevelopmental syndrome^33^, and non-coding intronic variants are amongst the strongest genetic risk factors for the neurodegenerative conditions amyotrophic lateral sclerosis (ALS) and frontotemporal dementia (FTD)^34, 35^.

Alongside changes in its function, the arrangement of Munc13-1 molecules at the active zone has been deemed critical for shaping synaptic transmission properties. In *D. melanogaster* neuromuscular junction synapses, the Munc13-1 ortholog Unc13a forms rings ∼70 nm around voltage-gated Ca^2+^ channels, contributing to the strong temporal coupling of synaptic transmission and Ca^2+^ influx triggered by an action potential^36, 37^. Upon synapse silencing, Unc13a expression is altered, with results pointing to an increase in the expression levels of Unc13a^38^ and/or compaction of the already-available Unc13a^39^, both associated with a homeostatic increase in synaptic strength. In mammalian synapses, Munc13-1 is arranged in nanoclusters and the number of nanoclusters is positively correlated with the strength of glutamate release^40^. CryoEM analysis and models based on the crystal structure of Munc13-1 promote the view that Munc13-1 forms hexameric rings surrounding one synaptic vesicle, acting in a cooperative manner to drive fusion^41, 42^. These rings have not been resolved in synapses yet, and, in general, tools are still lacking to visualize dynamic changes of Munc13-1 nanoarchitecture in mammalian synapses.

Here, we present a novel CRISPR/Cas9 knock-in mouse line, where we inserted a SNAP tag cassette at the endogenous *Unc13a* gene sequence to generate a Munc13-1 variant that is C-terminally fused to a SNAP tag. We validate the Unc13a^SNAP^ mouse line, and interrogate neurons via confocal, stimulated emission-depletion (STED) microscopy, and live cell imaging at confocal and super-resolution. Due to the complex nature of cultured neurons and brain tissue used, and because Munc13-1 is a protein with a moderate to low expression level^43^, we evaluated the performance of several SNAP tag substrates for efficient labeling.

We tested the bright and stable far-red SNAP dyes BG-JF_646_^44^ and BG-SiR-d12^45^, and, in addition, their membrane impermeable variants, SBG-JF_646_^13^ and SBG-SiR-d12, the synthesis and characterization of which we present here. Munc13-1 is a cytosolic protein and should not be labeled by membrane impermeable dyes, but given that some of our experiments were dealing with fixed and permeabilized preparations, we hypothesized that the charge originally installed to prevent membrane permeability may be useful in reducing background levels by repulsion. We report here that SBG-SiR-d12 successfully labels fixed cultured neurons and brain slices in terms of brightness and specificity and provided the best signal to noise ratio in many applications. Labeling using SBG-JF_646_, however, produced a substantial degree of background. We conclude that repurposing SBG-SiR-d12 for staining in permeabilized preparations is a promising approach to enhance labeling quality in complex samples, in cases where membrane permeability is irrelevant. We also conclude that the Unc13a^SNAP^ mouse line enables the detection of synapses and active zones at multiple scales, in live and fixed neurons, and thus may be used to characterize their rapid dynamics and plasticity *in vivo* or *in vitro*.

## RESULTS

### Generation and validation of an Unc13a-SNAP knock-in mouse line

We opted to generate a mouse model where endogenous Munc13-1 is C-terminally fused with a SNAP tag, with the two protein modules separated by a short and flexible, 9 amino acid long linker (sequence: (GGS)_3_, Figure 1A). Supporting our design strategy, we relied on a previously-generated knock-in mouse line where an enhanced yellow fluorescent protein (YFP) was added C-terminally to the Unc13a sequence^46^. A structural model of Munc13-1-SNAP generated by the recently-developed AlphaFold3^47^ (Figure 1B) predicts three important features, i.e. 1) the addition of the SNAP tag likely does not change the Munc13-1 structure, 2) the SNAP tag likely does not exhibit protein-protein interactions with Munc13-1, and, importantly, 3) the C-terminal C2C domain likely remains structured and accessible to protein-protein or protein-lipid interactions, both of which are critical for Munc13-1 function in SV priming and thus for setting the strength of neurotransmission^48–50^.

**Figure 1:**
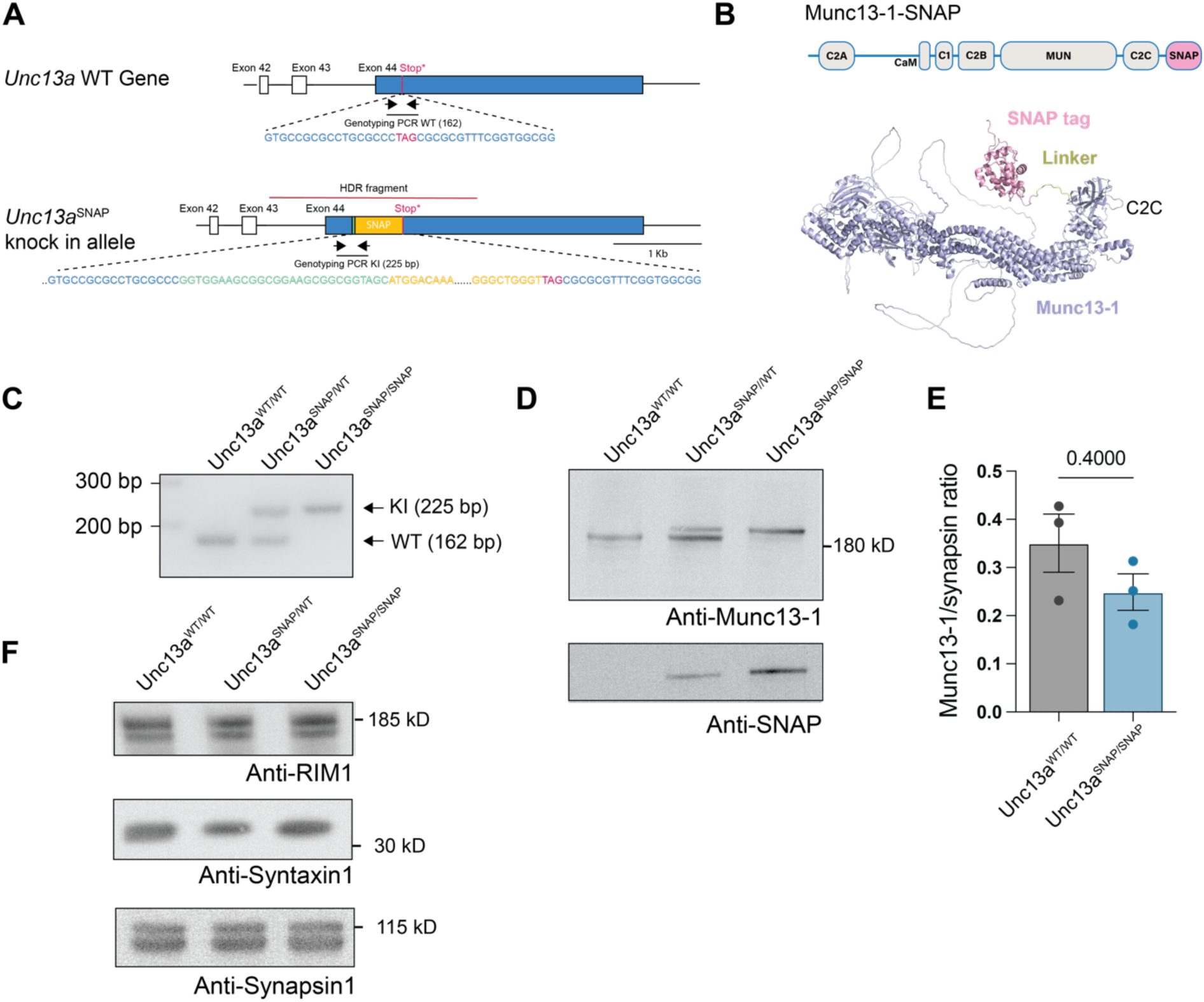
Validation of the Unc13a^SNAP^ mouse line. **(A)** Scheme of the 3’ region of the *Unc13a* gene in WT and in targeted, Unc13a^SNAP^ mice, including the DNA sequences at the site of SNAP tag insertion. Black arrows illustrate the location of the genotyping oligonucleotides, a black line illustrates the PCR fragment. The linker, stop codon, and genotyping oligonucleotides are not drawn to scale. **(B)** An illustration of the Munc13-1 domain structure in the Unc13a^SNAP^ mice (up) and AlphaFold3 structural prediction of the Munc13-1-SNAP protein (down). **(C)** Genotyping PCR results for the indicated genotypes. **(D)** A Western blot analysis of Munc13-1 (upper blot) and the SNAP tag (lower blot) in brain synaptosome homogenates from WT, heterozygous and homozygous Unc13a^SNAP^ mice. **(E)** Quantification of Munc13-1 levels from samples as in **(D)**. Data represents mean ± SEM from three independent experiments, Mann-Whitney test for statistical significance. **(F)** A Western blot analysis of RIM1, Syntaxin 1 and Synapsin 1 in brain homogenates from WT, heterozygous and homozygous Unc13a^SNAP^ mice.

The Unc13a^WT^ allele (Figure 1A) was targeted by site-directed CRISPR-Cas9 mutagenesis. The correct integration of the SNAP tag sequence was validated by long-range location PCRs (see Materials and Methods and the Supporting Information section for more information). In subsequent PCR analysis in genomic DNA from wild-type (WT), heterozygous (Unc13a^WT/SNAP^), and homozygous (Unc13a^SNAP/SNAP^) knock-in mice, we were able to amplify a DNA fragment spanning the last *Unc13a* exon and the SNAP tag cassette sequence (Unc13a^SNAP^ allele), confirming correct integration and enabling routine mouse genotyping (Figure 1C and Supporting Information section).

The resulting homozygous Unc13a^SNAP^ mice (Unc13a^em1(SNAP)Bros^) were viable and fertile, and exhibited no observable changes in survival, breeding performance, or cage behavior. Monitoring the well-being of the mice to adulthood (up to 8 months of age), we did not observe burden inflicted by the genetic modification. Because Munc13-1 loss results in perinatal lethality shortly after birth^26^, we conclude that the Munc13-1-SNAP protein fusion is functional.

Next, we evaluated the expression levels of Munc13-1 by Western blot analysis of a crude synaptosomal fraction (P2) from WT, heterozygous, and homozygous mouse brains. We found a non-significant, mild reduction in the expression levels of Munc13-1-SNAP in comparison to the Munc13-1 WT protein (Figure 1D, 1E). Interestingly, in samples from heterozygous mice the expression levels of the WT Munc13-1 protein appeared higher than that of the tagged Munc13-1 (Figure 1D, middle lane), which may indicate slightly reduced stability of the tagged protein in the presence of the WT form. This phenomenon has also been reported for the Munc13-1-eYFP fusion protein^46^, and should be considered when working with heterozygous mice. To confirm that the Munc13-1-SNAP fusion protein is fully functional, we conducted an electrophysiological analysis of synaptic transmission in glutamatergic excitatory neurons obtained from WT or Unc13a^SNAP^ littermate brains. We did not observe statistically-significant changes in the pattern or in the magnitude of synaptic transmission parameters, with the exception of a statistically-significant increase in the frequency of miniature excitatory postsynaptic currents frequency (Supporting Information, Figure S1), and conclude that the genetic integration of a SNAP tag at the C-terminus of Munc13-1 does not lead to overt changes in Munc13-1 function or expression.

Next, we established the expression of the SNAP tag cassette in samples from heterozygous and homozygous Unc13a^SNAP^ mice (Figure 1D), and excluded the presence of truncated Munc13-1-SNAP protein fragments, highlighting the stability of the tagged protein variant (Supporting Information, Figure S2). Finally, we found no change in the expression level of the major Munc13-1 interacting proteins syntaxin 1A/B and RIM1 (Regulating synaptic membrane exocytosis protein 1), as well as in the levels of Synapsin 1 in synaptosomal fractions from Unc13a^SNAP/SNAP^ mouse brains and WT littermate samples (Figure 1F).

To provide additional evidence to support the proper expression of Munc13-1-SNAP in Unc13a^SNAP^ neurons, we performed immunocytochemical analysis in hippocampal neuronal cultures from littermate WT and Unc13a^SNAP/SNAP^ mouse brains (Figure 2A, 2B, Table 1). We immunolabeled the neurons with 1) an antibody to label the Munc13-1 protein, 2) an antibody against Bassoon, a presynaptic active zone marker, and 3) an antibody against MAP2 to label the dendritic extensions of the neuron. We quantified the expression levels of Munc13-1 in WT and in Unc13a^SNAP/SNAP^ samples based on the Munc13-1 antibody signal intensity within regions of interest defined by the Bassoon signal, and found no changes between samples (Figure 2C). We also quantified the intensity of the signal arising from the labeling of Bassoon, and found no differences between the two genotypes (Figure 2D), indicating that the expression of Munc13-1-SNAP does not change Bassoon expression. To evaluate whether Munc13-1-SNAP is correctly localized in the active zone, we determined the colocalization coefficient between the Bassoon and Munc13-1 signals (Figure 2E). We found a high degree of colocalization (mean correlation coefficient values, WT, 0.545 ± 0.014, n=55 images, Unc13a^SNAP/SNAP^, 0.53 ± 0.017, n=54 images, Mann-Whitney test, see Supporting Information Figure S5 for data plotted for ROIs). We conclude that Munc13-1-SNAP is comparably-expressed and localized properly at presynaptic active zones.

**Figure 2:**
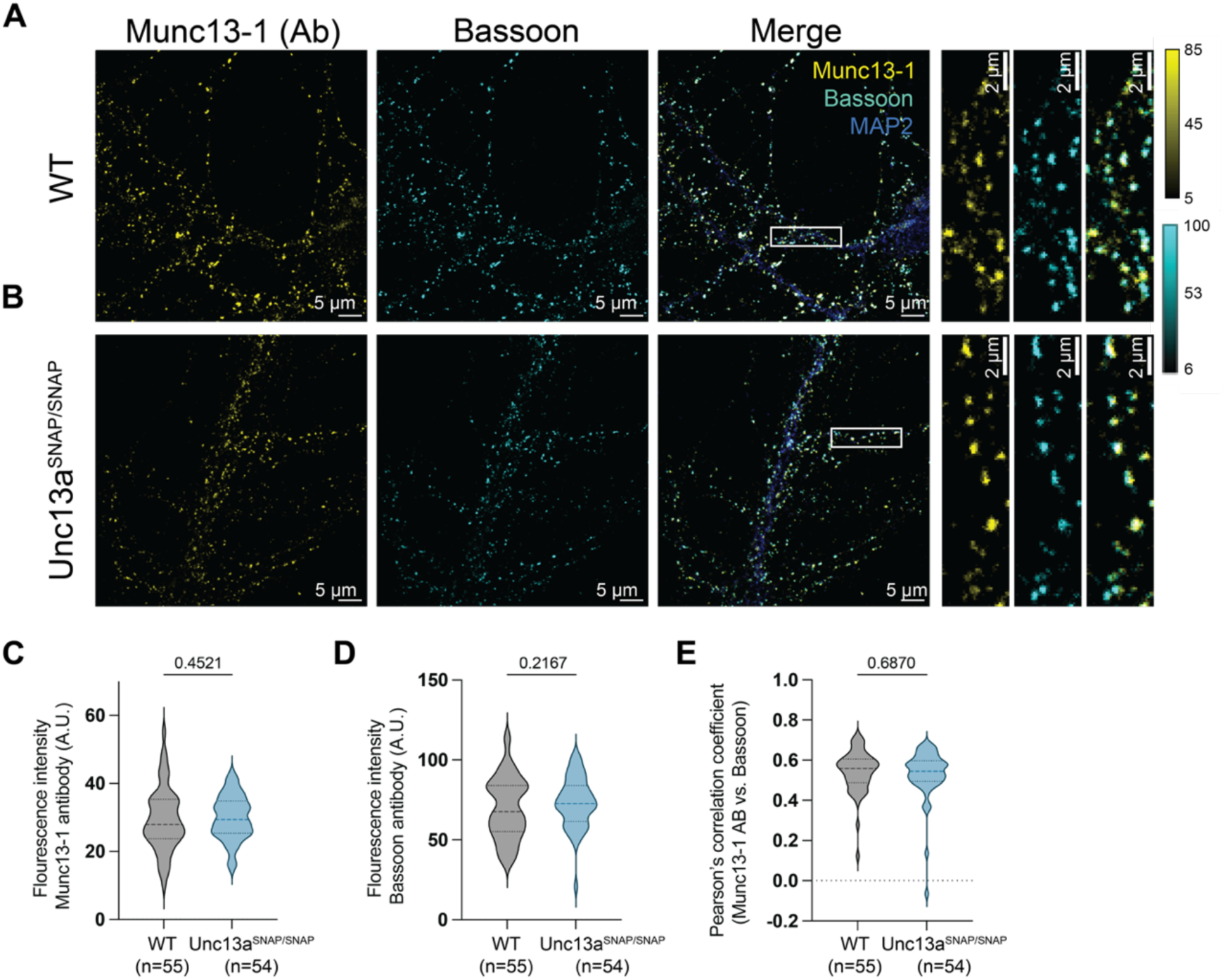
Validation of Munc13-1 localization in hippocampal neurons from Unc13a^SNAP/SNAP^ mice. Example images of cultured hippocampal neurons from WT **(A)** and Unc13a^SNAP/SNAP^ mouse brains **(B)**, immunolabeled with antibodies (Ab) against Munc13-1, the active zone marker Bassoon, and MAP2 (Scale bar: 5 µm, inset, 2 µm). On the right: magnification of the regions indicated by white boxes in the merged image. **(C)** Quantification of the fluorescence signal arising from antibody labeling of Munc13-1 and **(D)** of the fluorescence signal arising from antibody labeling of Bassoon in neurons from WT and Unc13a^SNAP/SNAP^ mice from images as in A and B. **(E)** The colocalization of Munc13-1 within Bassoon-labeled regions of interest, evaluated by calculating the Pearson’s correlation coefficient. In all violin plots, lines represent the median, 25% and 75% quartiles. Statistical significance was evaluated using a two-tailed Mann-Whitney test. n values represent the number of analyzed images, which were obtained from 3 independent experiment per condition (See Table 1 for a summary of the parameters used during microscopy experiments). A.U.: arbitrary units

### The development of SBG-SiR-d12

Following the validation of the knock-in mouse line, we next tested the efficiency of Munc13-1 labeling *via* the SNAP tag in fixed cultured primary neurons. In selecting the SNAP-dyes we opted for bright, stable, and far-red dyes. Silicon Rhodamine (SiR), and the next generation fluorophores JF_646_ and SiR-d12, were developed to exhibit boosted brightness without loss of resolution^44, 45^. Conjugated to BG (i.e. BG-JF_646_ and BG-SiR-d12), these dyes can be used for SNAP tag labeling. To improve the specificity of staining, sulfonated BG (SBG) substrates have been created^13^ and benchmarked in complex tissue^51^. The SBG moiety was originally designed for extracellular protein labeling, as it renders the conjugates membrane-impermeable. Thus, SBG-dye conjugates are not anticipated to label Munc13-1-SNAP, which is a cytosolic protein. Nonetheless, with the knowledge that the SBG moiety renders the conjugate less lipophilic, thus improving water solubility over prolonged periods of time^10^, we considered that the charged SBG-dye conjugate might reduce background levels through surface repulsion and generate less non-specific deposits. Cytosolic proteins could still be labeled, if the sample has been fixed and permeabilized prior to the SNAP dye application. We report here the synthesis and validation of a novel membrane impermeable SNAP dye, SBG-SiR-d12 (see methods and Supporting Information section). We were able to obtain and validate SBG-SiR-d12 against a cohesive palette of far-red dyes for SNAP tag labeling (Figure 3A).

**Figure 3:**
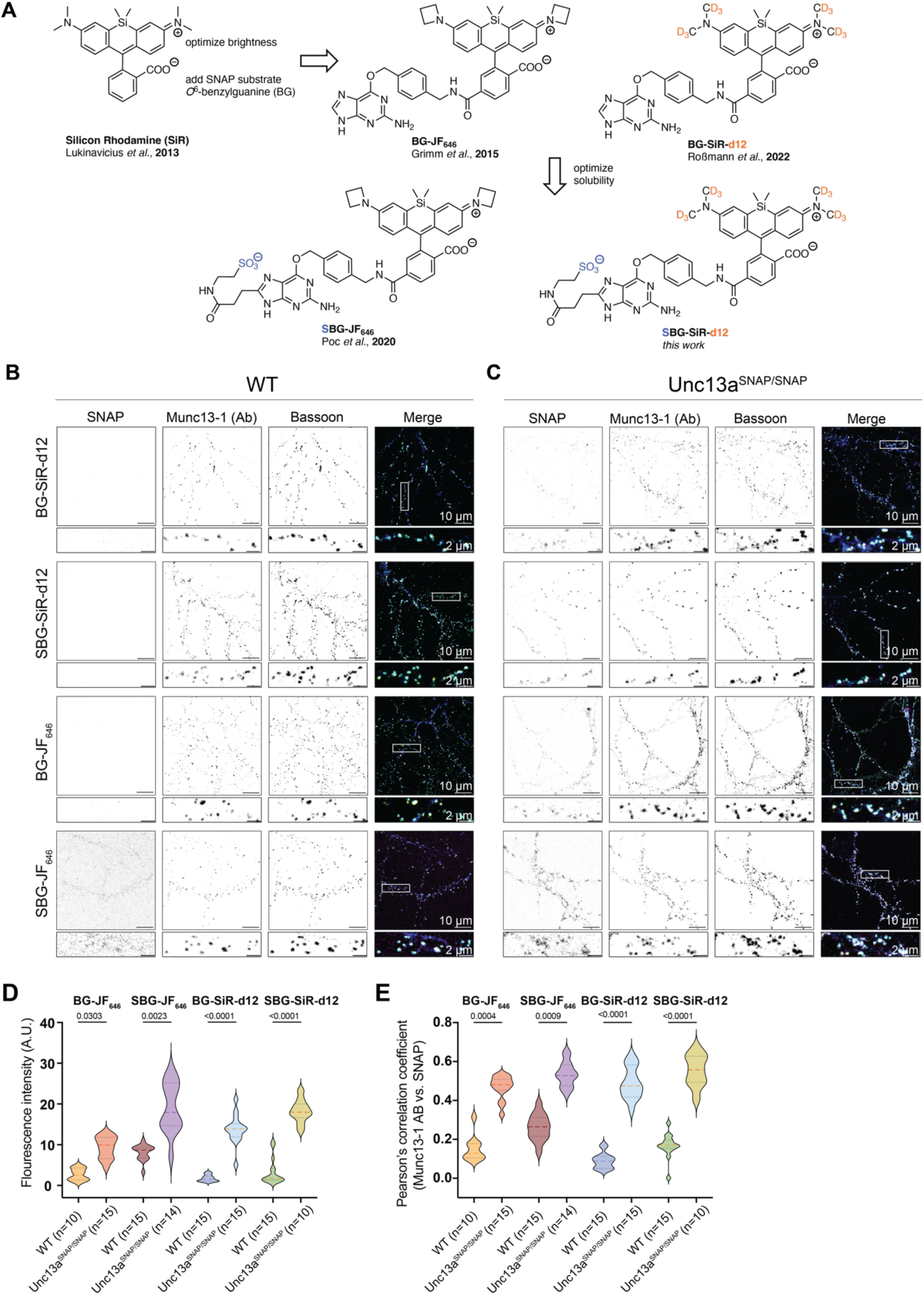
Development of SBG-SiR-d12 and validation of the SNAP tag functionality in cultured, fixed hippocampal neurons. (**A**) Chemical structures of silicone rhodamine variants for SNAP tag labeling, including SBG-SiR-d12 developed here. (**B-C**) Example images of fixed cultured hippocampal neurons from WT (**B**) and Unc13a^SNAP/SNAP^ (**C**) mouse brains, immunolabeled with the SNAP tag compounds BG-JF_646_, SBG-JF_646_, BG-SiR-d12, SBG-SiR-d12, as well as with antibodies (Ab) against Munc13-1, Bassoon, and MAP2 (Scale bar: 10 µm, inset, 2 µm). The location of the magnified regions below each image are indicated by the white box in the merged image. (**D**) Quantification of the fluorescence signal arising from Munc13-1-SNAP labeling by BG-JF_646_, SBG-JF_646_, BG-SiR-d12, SBG-SiR-d12, in neurons from WT and Unc13a^SNAP/SNAP^ mice. (**E**) Colocalization of signals arising from simultaneous Munc13-1 labeling by an anti-Munc13-1 antibody and via the SNAP tag, using the Person’s correlation coefficient. In the violin plots, lines represent the median, and 25% and 75% quartiles. In all figures, n represents the number of images analyzed, and a two-sided Kruskal-Wallis test followed by Dunn’s test for multiple comparisons was used to determine statistical significance. Data was obtained in three independent experiments (Table 1). A.U.: arbitrary units.

To test for the functionality and membrane (im)permeability of SBG-SiR-d12, HEK293 cells were transfected with a construct encoding for the fusion protein SNAP-TM-HTP, where an extracellular SNAP tag is separated by a transmembrane (TM) domain from an intracellular Halo Tag protein (HTP)^52^ (Supporting Information, Figure S3A). We then applied either BG-SiR-d12 (upper panel) and SBG-SiR-d12 (lower panel), together with chloroalkane-JF_519_ (CA-JF_519_), which is used to control for the expression of the SNAP-TM-HTP construct. Comparing BG-SiR-d12 and SBG-SiR-d12, we observed that the staining pattern of SBG-SiR-d12 is membrane-restricted, while BG-SiR-d12 also labels a cytosolic protein pool (e.g. membrane proteins still residing at the endoplasmic reticulum). This result suggests that SBG-SiR-d12 is less membrane permeable.

Next, we performed *in vitro* measurements to characterize pH sensitivity and tentative background staining behavior. We subjected 100 nM of SBG-SiR-d12 or SBG-JF_646_ to a pH titration (Supporting Information, Figure 3B) to gain information about their fluorogenicity. We found SBG-JF_646_ to be pH sensitive at more basic conditions while SBG-SIR-d12 was insensitive in a pH range from 4.2 to 9.2 (Supporting Information, Figure 3B). In a parallel experiment, we titrated BSA up to a concentration of 10 mg/mL, and found SBG-JF_646_ to give higher values in fluorescence polarization, indicating an increased tendency towards unspecific binding (Supporting Information, Figure 3C).

### Labeling of Munc13-1-SNAP in fixed samples using BG-dye compounds

We used primary neuronal cultures from WT and Unc13a^SNAP/SNAP^ mice, and tested BG-JF_646,_ SBG-JF_646_, BG-SiR-d12 and SBG-SiR-d12 for their efficiency in labeling Munc13-1-SNAP. We immunolabeled fixed and permeabilized neurons with antibodies against Munc13-1, Bassoon, and MAP2, and applied one of the BG-dye conjugates mentioned above (Figure 3B, 3C, Supporting Information, Table 1). Neurons from Unc13a^SNAP/SNAP^ mice were efficiently-labeled by all four SNAP-targeted dyes (Figure 3C, panel ‘SNAP’). In WT, negative control neurons, we did not observe unspecific labeling, except in the case of SBG-JF_646_, where substantial background was observed (Figure 3B, left panel ‘SNAP’). The unspecific stickiness (Supporting Information, Figure S3C) might be the cause for the less crisp performance of SBG-JF_646_ compared to SBG-SiR-d12, potentially also in combination with the pH sensitivity, for instance when unspecific binding occurs at basic protein microenvironments. We analyzed the signal intensity arising from the SNAP tag labeling (Figure 3D), and found that the fluorescent signal was strongest for SBG-JF_646_ and SBG-SiR-d12 in knock-in synapses. Importantly, the best signal to background ratio, which we define here as the fold-change in the median value of the signal intensity between Unc13a^SNAP/SNAP^ synapses and WT synapses for each SNAP dye, was superior for SBG-SiR-d12 (BG-JF_646,_ 3.8-fold, SBG-JF_646_, 2.05-fold, BG-SiR-d12, 8.4-fold, SBG-SiR-d12, 10.6-fold).

To demonstrate labeling specificity, we assessed the colocalization between signals arising from labeling of the same antigen: the signal for Munc13-1 arising from the SNAP tag labeling, and the signal of Munc13-1 arising from the antibody labeling. We found, as expected, a high degree of colocalization for all dyes tested (Figure 3E). We conclude that the SNAP tag conjugated to Munc13-1 is functional, and enables specific and bright labeling of Munc13-1 in fixed tissue at confocal resolution. SBG-SiR-d12 has risen as the dye of choice to label the SNAP tag in permeabilized, fixed samples.

We then tested whether the Unc13a^SNAP^ mice can be useful in imaging presynaptic terminals in slices of fixed brain tissue, which are typically difficult to label due to a high degree of non-specific binding of antibodies and reduced antibody permeability (Figure 4, Supporting Information Figure S4, Table 1). Sagittal sections from Unc13a^WT^ and Unc13a^SNAP/SNAP^ mice were labeled with either BG-JF_646_, SBG-JF_646_, BG-SiR-d12 and SBG-SiR-d12, in parallel to immunolabeling with an antibody against the presynaptic marker Synaptophysin, and a nuclear stain (DAPI). We obtained labeling with SBG-SiR-d12 in regions of high synapse density and where Munc13-1 has been shown to be enriched in a Munc13-1-YFP knock-in mouse line^46^, for example in the cortex, in the hippocampal *Stratum Orients* and *Stratum Radium*, and in the molecular layer of the cerebellum formation. This signal colocalized with antibody-stained Synaptophysin 1 signal, and little background was evident in WT control sections. The signal to background ratio of intensity measured in slices obtained from Unc13a^SNAP/SNAP^ and WT mice was 2.4 for SBG-SiR-d12, and lower for other dyes tested (BG-JF_646,_ 1.8-fold, SBG-JF_646_, 1.3-fold, BG-SiR-d12, 0.96-fold, Figure S4). We conclude that fixed tissue from Unc13a^SNAP^ mouse brains can be used for the rapid labeling of presynaptic terminals.

**Figure 4:**
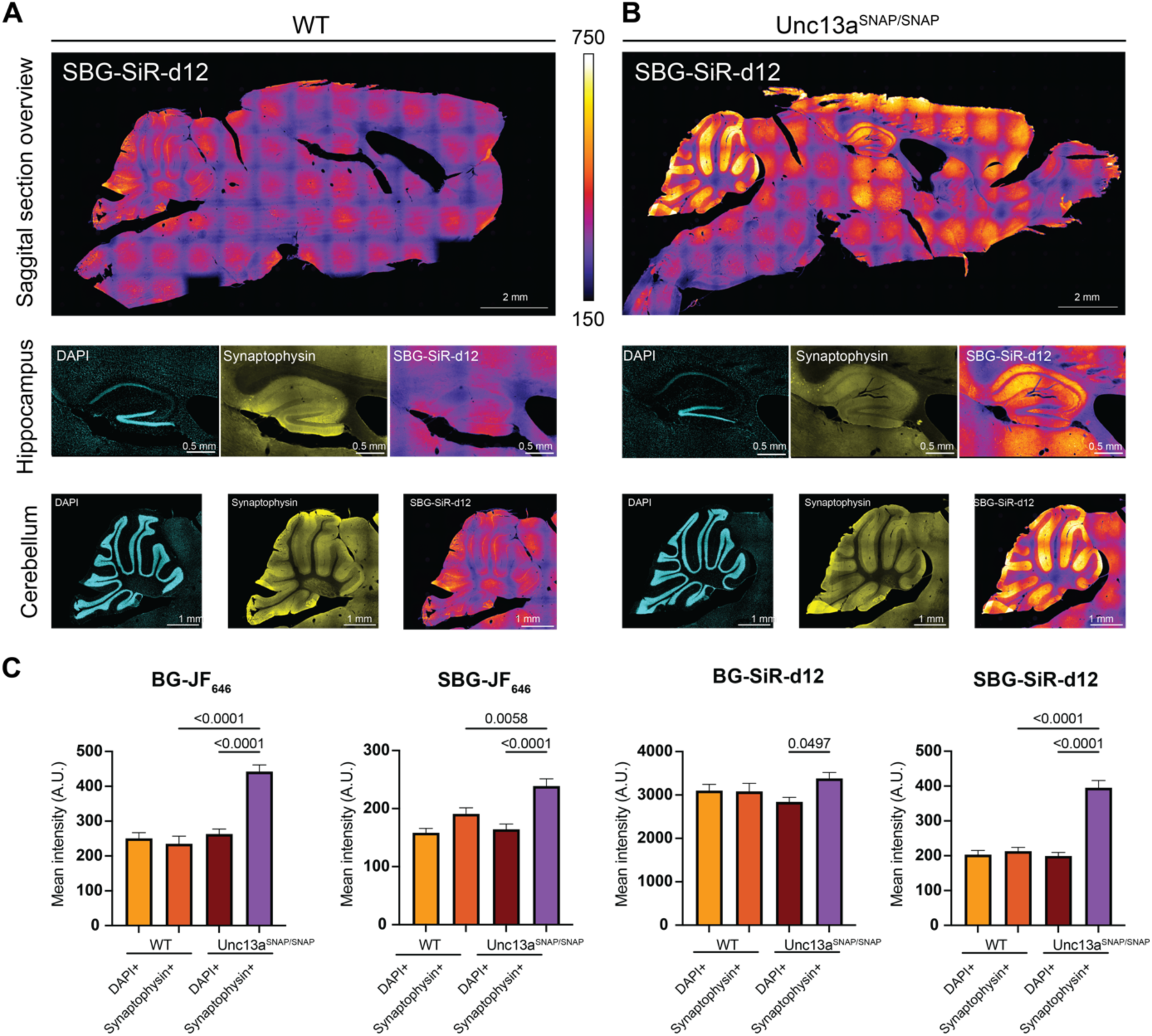
Labeling of presynaptic terminal in brain sections of Unc13aSNAP/SNAP mice. Sagittal brain sections from WT (**A**) and Unc13a^SNAP/SNAP^ (**B**) mice were stained with 1 µM SBG-SiR-d12, an antibody against Synaptophysin 1 (yellow, to stain synapses), and with DAPI (cyan), to stain cell nuclei. Example images in the region of the hippocampus (middle) and of the cerebellum (down). (**C**) Quantification of SNAP labeling overlapping with DAPI or Synaptophysin 1 signal in the cerebellum. The mean intensity of the SNAP signal was measured in 10 randomly-selected ROIs containing positive signals for either DAPI or Synaptophysin 1 (N=2 slices per condition, 20 ROIs for each column and genotype, A.U.: arbitrary units). Mann-Whitney test for statistical significance.

### Super-resolution imaging of Munc13-1-SNAP

Considering the ample data indicating that changes in Munc13-1 nano-organization can underline synaptic plasticity^36, 37, 40^, we were interested in establishing whether Munc13-1-SNAP can be imaged at super-resolution, to enable the localization of Munc13-1 beyond the diffraction limit^10^. We used stimulated emission depletion (STED) nanoscopy and imaged synapses from WT and Unc13a^SNAP/SNAP^ hippocampal neurons in culture, fixed and stained with antibodies against MAP2 (dendrites), Munc13-1, Bassoon, and one of the SNAP-targeting dyes described above. Example images of three synapses per compound are presented in Figure 5A (WT) and 5B (Munc13-1-SNAP) (See also Table 1). Using antibody staining, we identified Munc13-1 puncta that were localized in proximity to the active zone protein Bassoon, and were in part concentrated in nanoclusters. Imaging of Munc13-1 via the SNAP tag resulted in specific signals in Unc13a^SNAP/SNAP^ synapses. As in confocal imaging, we found a high degree of background when labeling WT samples with SBG-JF_646_, and the best signal to background ratio when using SBG-SiR-d12 (Figure 5C, BG-JF_646,_ 4.9-fold, SBG-JF_646_, 1.8-fold, BG-SiR-d12, 7.2-fold, SBG-SiR-d12, 9.6-fold). Signal intensities, however, were weaker compared to the antibody staining, which is expected due to the lack of signal amplification by primary and secondary antibodies. The SNAP tag signals were colocalized with the signals arising from the Munc13-1 antibody labeling (Figure 5D).

**Figure 5:**
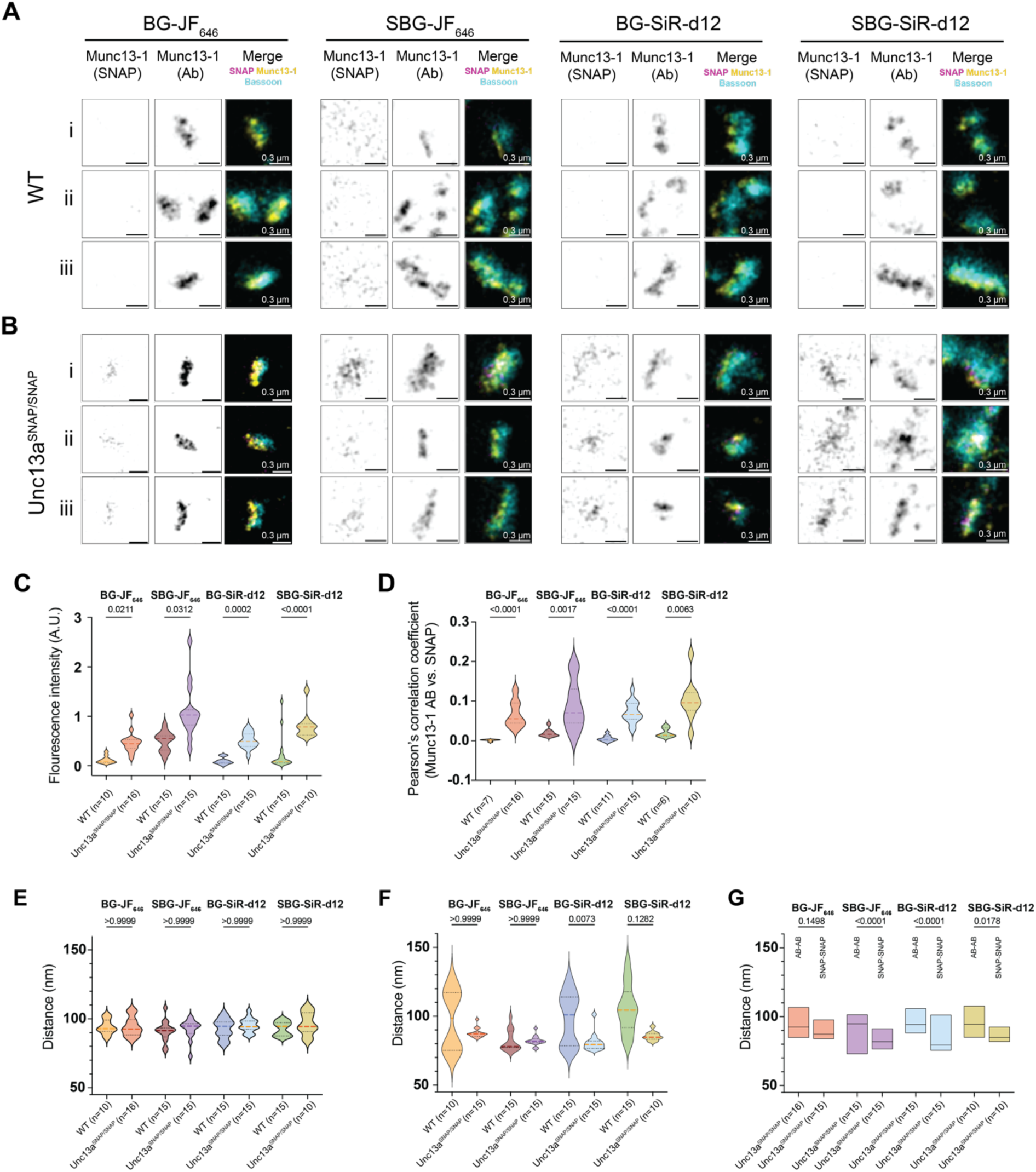
STED microscopy of Munc13-1-SNAP. **(A)** Example images of synapses from WT and **(B)** Unc13a^SNAP/SNAP^ neurons labeled with the SNAP tag compounds BG-JF_646_, SBG-JF_646_, BG-SiR-d12, SBG-SiR-d12, as well as with antibodies (Ab) against Munc13-1 and Bassoon, and imaged using STED microscopy. Three synapses are illustrated per condition (Scale bar: 0.3 µm) **(C)** Quantification of the fluorescence signal arising from Munc13-1-SNAP labeling by BG-JF_646_, SBG-JF_646_, BG-SiR-d12, SBG-SiR-d12, in neurons from WT and Unc13a^SNAP/SNAP^mice. **(D)** Colocalization of signals arising from Munc13-1 antibody labeling and SNAP tag labeling using the Pearson’s correlation coefficient. **(F-H)** Munc13-1 puncta-to-puncta distance measured based on **(F)** antibody labeling or **(G)** SNAP tag labeling, and **(H)** a comparison of antibody labeling versus SNAP labeling per SNAP dye tested. In the violin plots, lines represent the median, and 25% and 75% quartiles. In all figures, n represents the number of images analyzed, and a two-tailed Kruskal-Wallis test followed by Dunn’s test for multiple comparisons was used for the analysis of statistical significance. Data were obtained from three cultures. A.U.: arbitrary units.

We then used the images to analyze the distances between Munc13-1 puncta. No differences in puncta-to-puncta distances were observed when comparing WT and Unc13a^SNAP/SNAP^ synapses stained via an antibody against Munc13-1, indicating that Munc13-1 tagging does not disrupt the distribution of Munc13-1 (Figure 5E). The distances between Munc13-1 puncta obtained based on the SNAP tag staining were broadly distributed in WT synapses, consistent with the background nature of the signal (Figure 5F), whereas a tight distribution was measured in Unc13a^SNAP/SNAP^ neurons. Finally, we compared the puncta-to-puncta distances generated by SNAP tag staining with those measured using antibody staining. We found that distances were significantly shorter when labeled via the SNAP tag across all tested dyes (Figure 5G). These findings suggest that labeling of Munc13-1 via endogenous SNAP tagging may serve as a valuable complementary approach to antibody labeling.

### Live labeling and imaging of Munc13-1-SNAP

One significant advantage of self-labeling protein tags is that they offer the possibility to monitor proteins in living cells. We therefore tested whether we could utilize the Unc13a^SNAP^ neurons for monitoring Munc13-1 using live imaging. In our protocol, WT and Unc13a^SNAP/SNAP^ neurons were incubated with 0.1 µM BG-SiR-d12 or BG-JF_646_ dissolved in a physiological buffer for 45-60 min, and were then allowed to recover for 24-48 hours in growth media before subjected to imaging (Figure 6A). The half-life of Munc13-1 has been estimated in neuronal cultures to be in the range of 3.5-4 days^53, 54^, which is compatible with such long recovery times. Very short recovery times resulted in a significant level of background staining, but we did not test intermediate recovery times. The neurons were then subjected to live cell imaging on a STED microscope with 640 nm Excitation and 775 nm depletion at 37 °C. In confocal mode, we observed strong punctate staining that was highly-specific for Unc13a^SNAP/SNAP^ neurons, and merely absent in WT neurons (Figure 6B). We then acquired images at STED resolution, and readily observed Munc13-1 nanoclusters (Figure 6C). To ascertain that these signals largely represents synapses, we conducted a second experiment where we used lentiviral-mediated transduction to sparsely express a GFP-tagged vesicular glutamate transporter (VGLUT1), a presynaptic marker, in Unc13a^SNAP/SNAP^ neurons (Figure 6D). We observed GFP and BG-SiR-d12 puncta in close apposition. Subjecting the images to STED imaging, we again observed Munc13-1 clusters colocalizing with the GFP label (Figure 6D lower and right panel). Finally, we quantified the UNC13A nanocluster diameter, and found a median diameter of 135-160 nm. This diameter is larger than that described by others^40^, reflecting the limited labelling density and an estimated resolution of 60 nm for live STED (see comment in figure 6E as well). We conclude that the Munc13-1-SNAP mice enable stable and bright live imaging of presynaptic terminals at confocal- and super-resolution.

**Figure 6:**
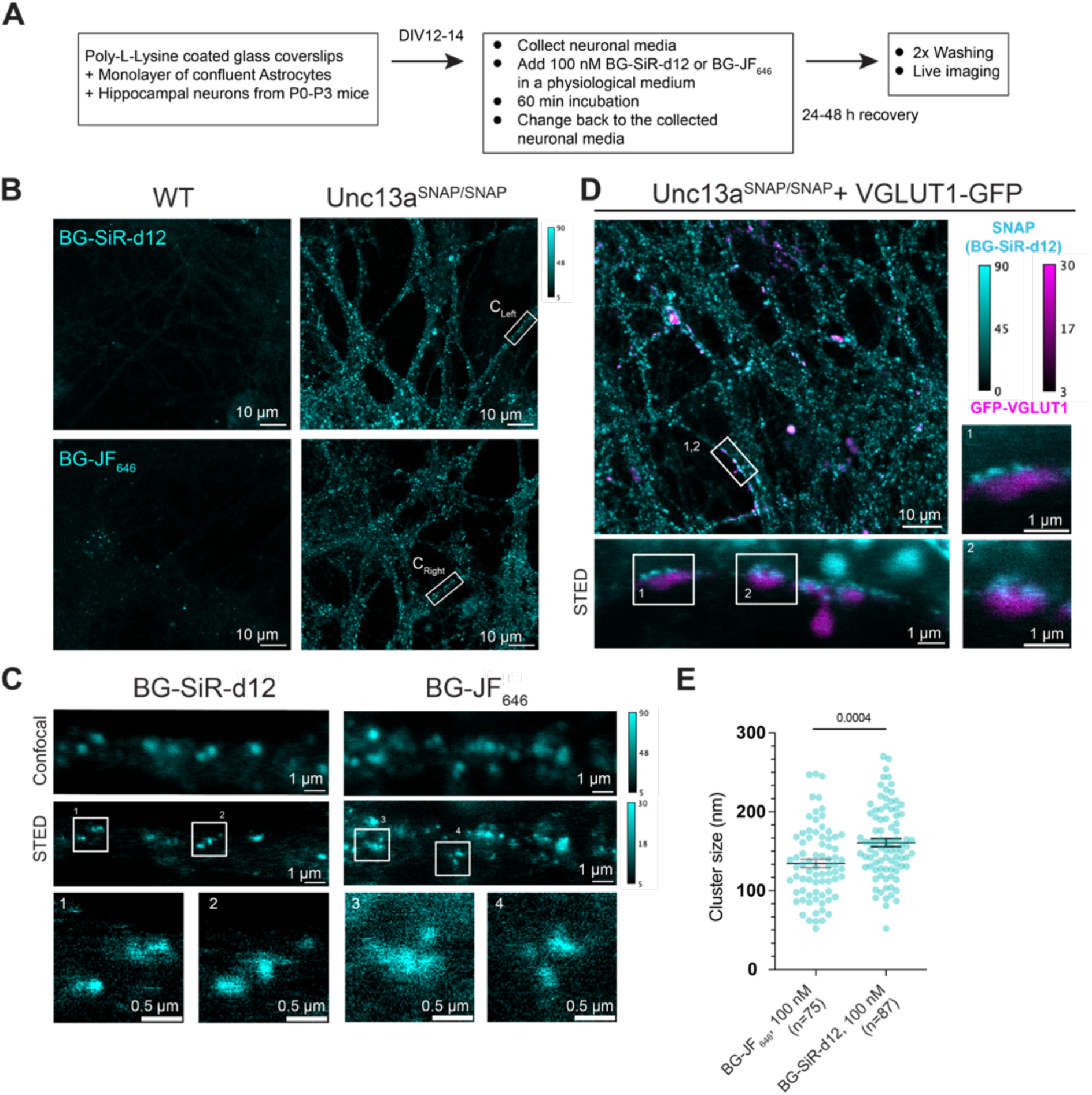
Live imaging of cultured neurons from Unc13a^SNAP/SNAP^ mice. (**A**) Labeling protocol established for live imaging of Munc13-1-SNAP and using a Nikon Eclipse TI microscope with a STEDYCON system. (**B**) Example confocal images of live WT (left) and Unc13a^SNAP/SNAP^ neurons (right) stained with BG-SiR-d12 (up) or BG-JF_646_ (down), and (**C**) STED imaging on selected processes from the images in (B). (**D**) Example images of Unc13a^SNAP/SNAP^ neurons that were infected with a lentivirus expressing a GFP-tagged vesicular glutamate transporter (GFP-VGLUT1, magenta), stained with BG-SiR-d12 (cyan) and imaged at confocal (up) and STED resolution (down and right). (**E**) Quantification of Munc13-1 nanocluster diameter using line scans and gaussian fit in images obtained during live STED imaging. In Figure 6E, data from individual puncta is plotted as mean ± SEM, Kruskal-Wallis test followed by a Dunn’s test for multiple comparisons.

## DISCUSSION

Changes in synaptic function have been repeatedly linked to changes in the organization and composition of the neurotransmitter release machinery. At the presynaptic compartment, the proximity (coupling) of release sites to voltage-gated Ca^2+^ channels determine the strength and probability of neurotransmitter release and can be dynamically modulated following plasticity-inducing triggers^55–58^. Plasticity also evokes changes in the size of the active zone^59^, and in the abundance of key synaptic proteins^36, 38, 60,53^. Munc13-1 is a central presynaptic protein that is absolutely essential for neurotransmitter release, and changes in Munc13-1 abundance and nanoarchitecture are emerging as critical for setting neurotransmitter release properties^40, 61^. However, tools are still lacking to monitor such nanoarchitectural changes for endogenous Munc13-1 *in vivo* and *in vitro*.

We used CRISPR-Cas9 to knock-in the SNAP tag cassette in the last exon and prior to the stop codon of the *Unc13a* gene. We used the SNAP cassette to complement already-available tools for synapse nanoscopy, in particular the PSD95-HaloTag line^62^ that has proven as highly useful for analyzing postsynaptic density at super-resolution^63^, but also for imaging synapses in brain slices^62^. Successful targeting in mouse embryos enabled us to establish a mouse line constitutively expressing Munc13-1-SNAP at endogenous expression levels. In hippocampal neuronal cultures of this mouse, Munc13-1-SNAP is expressed at WT levels and localizes to the active zone. Fixed neuronal culture samples and brain slices could be labeled via the SNAP tag ligands following fixation and were successfully imaged via confocal and/or STED imaging. Importantly, we demonstrated efficient and specific labeling of live neurons in culture, which opens the door for studies where labeling Munc13-1 at a timepoint of choice and monitoring its dynamics in living cells are possible. Together, we validate the Unc13a^SNAP^ mice for future studies of synapse and active zone nanoarchitecture.

The key advantages of self-labeling tags include great flexibility in dye selection and a large and consistently growing toolbox of chemicals available. Importantly, reagents do not only differ in the wavelength of the fluorophore attached to them, but also in their chemical and biological properties, e.g. they undergo uptake to different extents, and have variable solubilities. Nonetheless, a large fraction of reported investigations using self-labeling tags rely on the application of one derivative only. We show here by testing several BG derivatives simultaneously that the performance of different BG-fluorophore compounds is assay-dependent and needs to be carefully controlled^3, 64^. Moreover, we add SBG-SiR-d12 as an additional dye in the SNAP dye toolbox. While a priori, using a sulfonated and impermeable dye seems counterintuitive, we speculated that SBG dyes will be functional in the context of fixed and permeabilized samples. SBG-SiR-d12 outperformed all other conjugates in signal strength, and had the highest signal to background ratio in fixed cell (10-fold) and in complex brain tissue samples (2.4-fold). Its enhanced performance with respect to JF dyes could be attributed to less pH sensitivity and a smaller degree of unspecific binding, which either or in combination may result in more background signals stemming from SBG-JF_646_. We propose SNAP labeling in fixed cell cultures and slices with SBG-SiR-d12 as an alternative to membrane-permeable SNAP dyes. The protocol presented here joins protocols for SNAP tag labeling in the live setting, e.g. in immortalized cell lines^3^, *Drosophila* brain^65, 66^, in mice for neural identification and ablation^67^, for receptor localization including optical manipulation^68^, and in pancreatic islets^69^.

Further developments of imaging configurations, SNAP tag reagents and labeling protocols are expected to expand the range of experimentation possible using the Unc13a^SNAP^ mice^70–74^. The genetic fusion via CRISPR/Cas9 ensures the visualization of Munc13-1 at endogenous expression patterns and levels, independent of constraints related to antibody species, and in the context of neuronal networks in either fixed or live preparations. These key features will surely make the Unc13a^SNAP^ mice useful in the discovery of synaptic nanoarchitectural principles in the future.

## MATERIALS AND METHODS

### General

BG-JF_549_, SBG-JF_549_, and BG-SiR-d12 have been synthesized and used previously in our laboratories^10, 42^.

### Chemical Synthesis

SBG-SiR-d12 was synthesized as follows: In an Eppendorf tube 1.2 mg (2.0 µmol, 1.0 equiv.) of SiR-d12-COOH^45^ was dissolved in 100 μL/mg DMF and 8.0 equiv. of DIPEA. Upon addition of 2.3 equiv. TSTU (from a 41 mg/mL stock in DMSO) the reaction mixture was vortexed and allowed to incubate for 10 min, before 1.2 equiv. of SBG-NH_2_ was added. The mixture was vortexed again and allowed to incubate for 60 min before it was quenched by addition of 20 equiv. of acetic acid and 25 vol% of water. C18 RP-HPLC (MeCN:H_2_O+0.1% TFA = 10:90 to 90:10 over 45 minutes) provided the desired compound, SBG-SiR-d12, which was obtained as a blue powder after lyophilization in 42% yield (0.84 µmol) and aliquoted to 5 nmol to be stored at –20 °C.

### ^1^H NMR (600 MHz, MeOD-d_4_)

δ [ppm] = 9.29–9.21 (m, 1H), 8.23–8.16 (m, 1H), 8.12–8.08 (m, 1H), 7.71 (s, 1H), 7.50 (d, *J* = 7.9 Hz, 2H), 7.39 (d, *J* = 7.9 Hz, 2H), 7.20 (s, 2H), 6.86 (d, *J* = 9.1 Hz, 2H), 6.70 (dd, *J* = 9.3, 3.0 Hz, 2H), 5.59 (s, 2H), 4.59–4.55 (m, 2H), 3.54 (t, *J* = 6.5 Hz, 2H), 3.45–3.41 (m, 1H), 3.20–3.18 (m, 1H), 3.12 (t, *J* = 7.0 Hz, 2H), 2.90 (t, *J* = 6.5 Hz, 2H), 2.70 (t, *J* = 7.0 Hz, 2H), 0.63 (s, 3H), 0.57 (s, 3H).

### HRMS (ESI)

calc. for C_45_H_38_D_12_N_9_O_8_SSi [M+H]^+^: 916.4020, found: 916.4165.

Raw fluorescence and fluorescence polarization measurements were performed on a TECAN Spark Cyto (λ_Ex_ = 610±20 nm, λ_Em_ = 650±9 nm) and on a TECAN GENios Pro plate reader (λ_Ex_ = 610±15 nm, λ_Em_ = 650±20 nm), respectively. Stocks of substrates (100 nm) were prepared in PBS with either adjusted pH or additional BSA as indicated in a Greiner black flat bottom 96 well plate. Experiments were run in replicates and plotted in Prism 10.

### Animal study approval

The generation and use of the Unc13a-SNAP (Unc13a^em1(SNAP)Bros^) knockin mice were approved by the responsible local government organization (Niedersächsisches Landesamt für Verbraucherschutz und Lebensmittelsicherheit, 33.9-42502-04-13/1359 and 33.19-33.19-42502-04-20/3589). All the experiments performed in Berlin complied with European law and the state of Berlin animal welfare body (LAGeSo).

### Generation of an Unc13a-SNAP knock-in mouse line

Superovulated C57BL6/J females were mated with C57BL6/J males and fertilized eggs collected. In-house prepared CRISPR reagents (hCas9_mRNA, sgRNAs, preformed Cas9_sgRNA RNP complexes, and the dsDNA used as a repair template (HDR fragment), were microinjected into the pronucleus and the cytoplasm of zygotes at the pronuclear stage using an Eppendorf Femtojet. Importantly, all nucleotide-based CRISPR-Cas9 reagents (sgRNAs and hCAS9_mRNA) were used as RNA molecules and were not plasmid-encoded, reducing the probability of off-target effects, due to the short live of RNA-based reagents^75, 76^. The sgRNAs targeting the region around the Munc13-1 STOP-codon were selected using the guide RNA selection tool CRISPOR^77, 78^. The correct site-specific insertion of the HDR fragment was confirmed by two localization PCRs with primers upstream and downstream of the HDR sequence, followed by direct sequencing of the obtained PCR products. The sequences of the RNA and DNA fragments used to generate and validate the mice are available in the Supporting Information section. We term the line as Unc13a^em1(SNAP)Bros^. For questions about usage of the line, please contact Nils Brose and Noa Lipstein.

### Synaptosome preparation and Western blot analysis

A crude synaptosomal fraction (P2) was prepared as described in^79^ from mice age 7-11 weeks. The sample was solubilized to a final concentration of 2 mg/ml in 50 mM Tris/HCl pH 8, 150 mM NaCl, 1 mM CaCl_2_, 1 mM EGTA, 1% NP-40, 0.2 mM phenylmethylsulfonyl fluoride, 1 mg/ml aprotinin, and 0.5 mg/ml leupeptin, stirred on ice for at least 15 min at 4 °C, centrifuged at 4 °C, 20,000g for 5 minutes to remove insoluble material, mixed with Laemmli sample buffer and boiled for 10 minutes at 99 °C. 5 µg of each sample were loaded on a 4-12% Bis-Tris gel, and the proteins were separated and transferred to a nitrocellulose membrane. The membrane was washed three times with ddH_2_O, stained with Pierce™ Reversible Protein Stain Kit for Nitrocellulose Membranes (Thermo Scientific 24580) and scanned. Then, the membrane was blocked for 60 minutes with blocking buffer (5% (*w*/*v*) low fat milk in Tris buffered Saline (TBS) buffer supplemented with 0.1% Tween20 (TBS-T)), and blotted with the following antibodies diluted in blocking buffer for 2 hours at RT with gentle shaking: Rabbit anti Munc13-1 (Synaptic Systems 126 103, diluted 1:1000), Rabbit anti SNAP (New England Biolabs, P9310S, diluted 1:500), Mouse anti synapsin 1 (Synaptic Systems 106 011, diluted 1:1000), Mouse anti Syntaxin 1A/B (Synaptic Systems 110 011, diluted 1:2000), Rabbit anti Rim1 (Synaptic Systems 140 003, diluted 1:1000). The membrane was washed three times for 10 minutes each with TBS-T before blotted for 1 hour at RT with secondary antibodies conjugated with horseradish peroxidase (HRP) diluted in blocking buffer: Goat-anti-Rabbit-HRP (Jackson immune 111-035-114, diluted 1:5000) and Goat-anti-Mouse-HRP (Jackson immuno 115-035-146, diluted 1:5000). Before developing the chemiluminescence signal using the standard HRP signal amplification system, the membrane was washed three times for 10 minutes each in TBST buffer and once shortly in TBS buffer.

### HEK293T cultures

HEK293T cells were cultured in growth medium (DMEM, Glutamax, 4.5 g Glucose, 10% FCS, 1% PS, Invitrogen) at 37 °C and 5% CO_2_. 30,000 cells per well were seeded on 8-well μL slides (Ibidi) previously coated with 0.25 mg ml^−1^ poly-L-lysine (Aldrich, mol wt 70 000–150 000). The next day, 400 ng DNA was transfected using 0.8 μL Jet Prime reagent in 40 μL Jet Prime buffer (VWR) per well. Medium was exchanged against antibiotic-free media before the transfection mix was pipetted on the cells. After 4 hours incubation at 37°C and 5% CO_2_, medium was exchanged against growth media. After 24 hours cells were stained and imaged. All dyes were used at a concentration of 500 nM. 5 μM Hoechst 33342 was used to stain DNA. Staining was done in growth medium at 37 °C, 5% CO_2_ for 30 minutes. Afterwards cells were washed once in growth media and imaged live in cell imaging buffer (Invitrogen).

### Neuronal cultures

Primary hippocampal neurons were cultured as previously described^80^. Briefly, hippocampi were dissected from Unc13a^SNAP/SNAP^, Unc13a^WT/SNAP^, and WT/WT P0-P1 littermate mice, and were incubated at 37 °C gently shaking in a solution containing 0.2 mg/ml L-cysteine, 1 mM CaCl_2_, 0.5 mM ethylenediaminetetraacetic acid (EDTA) and 25 units/ml of papain (Worthington Biochemicals), pH 8 in Dulbecco’s modified eagle medium. After 45 minutes, the solution was replaced by a prewarmed solution containing 2.5 mg/ml Bovine Serum Albumin, 2.5 mg/ml trypsin inhibitor, 1% Fetal Bovine Serum, heat inactivated in Dulbecco’s modified eagle medium, and the hippocampi were incubated for 15 minutes at 37°C gently shaking. The solution was replaced by neuronal culture medium (Neurobasal™-A medium supplemented with 2% B-27^TM^ Plus Supplement, 1% GlutaMAX^TM^ supplement, and 0.2% Penicillin-Streptomycin), and the hippocampi were gently triturated to produce a cell suspension that was plated on PLL-coated coverslips for 2-3 weeks at 37 °C and 5% CO_2_. The culture contains primarily neurons and few astrocytes. Neurons were used for experiments at day-in-vitro 14-17.

### Lentiviral preparation

The construct encoding VGLUT1-GFP was a gift from C. Rosenmund and is based on the FUGW vector^81^, in which the ubiquitin promoter was exchanged by the human synapsin 1 promoter. Lentiviral preparation was carried out by the viral core facility of the Charite - Universitätsmedizin Berlin (vcf.charite.de) according to the protocol published in^81^ and modified as in^82^, using the helper plasmids provided by addgene # 8454 and # 8455^83^.

### Electrophysiology

Neurons were prepared from brains of P1 Unc13a^WT^ and Unc13a^SNAP/SNAP^ littermate mice and plated on WT astrocyte microisland cultures according to published protocols^80^, and kept at 37°C, 5% CO_2_ until recordings were made at DIV12-14. Whole-cell voltage-clamp data were acquired using a Axon Multiclamp 700B amplifier, Digidata 1440A data acquisition system, and pCLAMP 10 software (Molecular Devices). All recordings were performed using an external solution containing 140 mM NaCl, 2.4 mM KCl, 10 mM HEPES, 10 mM glucose, 4 mM CaCl_2_, and 4 mM MgCl_2_ (320 mOsm/l). The standard internal solution contained 136 mM KCl, 17.8 mM HEPES, 1 mM EGTA, 4.6 mM MgCl_2_, 4 mM NaATP, 0.3 mM Na_2_GTP, 15 mM creatine phosphate, and 5 U/ml phosphocreatine kinase (315–320 mOsm/l), pH 7.4. Recordings were made at room temperature (∼22°C). eEPSCs were evoked by depolarizing the cell from –70 to 0 mV for 1 ms duration. Basal eEPSCs were recorded at a frequency of 0.1 Hz. mEPSCs and mIPSCs were recorded for 50 s in the absence of tetrodotoxin. mEPSCs traces were filtered at 1 kHz and miniature events were identified using a sliding template function in Axograph or in IgorPro (5–200 pA, rise time 0.15–1.5 ms, half-width 0.5–5 ms). Analyses were performed using Axograph 1.4.3 (MolecularDevices) or Igor Pro (Wavemetrics). Electrophysiological data is presented as mean ± SEM.

### Immunocytochemical staining protocols

Experiments were made in neuronal cultures at days-in-vitro 14-17. Neurons were washed twice with PBS, and fixed by adding cold 1% paraformaldehyde for 10 minutes on ice. Neurons were again washed and leftover paraformaldehyde was quenched by adding 50 mM cold glycine for 10 minutes, followed by two additional washes with PBS. The cells were either stored in PBS at 4 °C or immediately used for immunostaining. The cell membrane was permeabilized with cold 0.25% Triton-X-100 in PBS for 10 minutes and rinsed once in pure PBS. To block nonspecific binding sites, the cells were incubated in cold 0.3% NGS in PBS for 20 minutes. All steps were performed with the multi-well plate stored on ice to prevent protein degradation.

To label Munc13-1 using the SNAP tag, 250 nmol BG-dye conjugates were diluted in blocking solution and applied to the neurons for 30 minutes at room temperature (RT) in the dark. Subsequently, the cells were extensively washed up to eight times in blocking solution and the following antibodies were applied in blocking buffer for immunolabeling: Chicken polyclonal MAP2 (Novus Biologicals NB300-213, diluted 1:1000), Rabbit polyclonal Munc13-1 (Synaptic Systems AB_126103, diluted 1:500), and Guinea Pig polyclonal Bassoon (Synaptic Systems AB_141004, diluted 1:500). Incubation was performed for 1 h at RT in the dark. To remove unbound antibodies, the cultures were washed three times with blocking solution before applying the following secondary antibodies conjugated with fluorophores: Anti-Chicken-405, Anti-Guinea Pig-488 and Anti-Rabbit-594 (all diluted 1:500). The neurons were incubated with the secondary antibodies for 30 minutes at RT in the dark and washed eight times in PBS to remove unbound products. The coverslips on which the neurons were fixed were mounted with ProLong™ Gold Antifade Mountant (Invitrogen™ P36934) on microscope glass slides and stored in the dark at RT for 48 hours until dry before imaging.

### Brain sections and immunohistochemistry

Animals were transcardially perfused with fixative (4% paraformaldehyde in 1X PBS, pH 7.4). Brains were post-fixed overnight at 4 °C, incubated in 30% sucrose in 1X PBS for a minimum of 48 h, and snap frozen with 2-methybutan on dry ice. Tissues were sectioned at 30 μm on a cryostat. Floating sections were kept in 30% ethylene glycol and 30% glycerol in 1X PBS at −20 °C. For immunostaining, sections were washed three times in 1X PBS before incubation with blocking solution containing 10% normal donkey serum (NDS) in TBS-T (0.05% Triton X-100 in 1X TBS). The slices were incubated with one of the SNAP tag dyes (1 µM) and anti-Synaptophysin1 (Synaptic Systems AB_101002, diluted 1:200) at 4 °C overnight in TBS-T containing 2% NDS. Sections were washed three times in 1X TBS-T, followed by incubation for 2 h at room temperature in the dark with secondary antibody (Alexa Fluor 488-AffiniPure Donkey Anti-Rabbit IgG, Jackson Research 711-545-152, diluted 1:1000,) and DAPI (Sigma MBD0015-1ML, diluted 1:10000). The sections were washed three times in 1X TBS-T before mounted onto microscope slides with ProLong Glass antifade (Invitrogen™P36982).

### Confocal and STED Imaging

Confocal and STED microscopy on fixed neuronal samples was performed using a Leica SP8 TCS STED FALCON (Leica Microsystems). Confocal images were collected using a time gated Hybrid detector (0.5–6 ns). Images of 1024×1024 pixel had a pixel size of 130 nm for confocal imaging and 20 nm for STED imaging. For imaging far red SNAP dyes, we used the following settings: excitation wavelength 646 nm, emission wavelength 656–751 nm. Brain sections were imaged on a Nikon Spinning Disk Field Scanning Confocal System using 10X objective (Plan Apo λD, NA 0.45), acquired in the “Large Image” mode of NIS 5.4 Software and stitched with 20% overlap between adjacent frames. For imaging DAPI, Synaptophysin 1 and SNAP dyes 405, 488 and 640 nm excitation lasers and 440/40, 525/50, 708/75 emission filters were used, respectively. CA-JF_519_ and SBG-SiR-d12 in live transfected HEK293Ts cells was imaged on a TIE Nikon epifluorescence microscope equipped with a pE4000 (cool LED), Penta Cube (AHF 66–615), 60 oil NA 1.49 (Apo TIRF Nikon) and a sCMOS camera (Prime 95B, Photometrics) operated by NIS Elements (Nikon). For excitation the following LED wavelengths were used: Hoechst: 405 nm, JF_519_: 480 nm, SiR-d12: 635 nm.

### Live Imaging

Live confocal and STED imaging was done on a STEDYCON system (Abberior Instruments GmbH, Göttingen), mounted on a Nikon Eclipse TI research microscope, equipped with a Plan APO Lambda 100X/1.45 NA oil objective (Nikon) and controlled by NIS Elements (Nikon). 24 h prior to live-cell imaging, an incubation chamber surrounding the microscope set-up was set to 37 °C. In order to provide stable focus during imaging the Perfect Focus System (Nikon) was used. Imaging was performed in a medium containing 140 mM NaCl, 2.4 mM KCl, 10 mM HEPES, 10 mM glucose, 2 mM CaCl2, and 2 mM MgCl2 (320 mOsm/l) at 37 °C. To capture a SNAP tag signal, excitation was evoked with a 640 nm diode laser and Emission was detected with a single counting avalanche photodiode (650-700 nm). The GFP-VGLUT1 signal was generated with excitation using a 488 nm diode laser. A pixel size of 200 nm x 200 nm or 20 nm × 20 nm were used in confocal and STED imaging, respectively. Confocal images were captured with a line accumulation of 1, while STED images utilized a line accumulation of 5.

### Image data analysis

Confocal and STED images (Figures 3, 5 and 6) were analyzed using routines generated in Matlab (The Mathworks Inc., Natick MA, USA, version R2022b). The Bassoon images were subjected to an automated thresholding procedure, to identify signals above background (using a threshold equal to the mean intensity value plus the standard deviation, calculated for the entire image). The resulting regions of interest (ROIs) were processed by an erosion procedure, using a kernel of 4 pixels, to remove noise events, and only retain synapse-like signals. The remaining ROIs were then automatically dilated, to include the surrounding synaptic areas, and the average intensity values were measured and reported, for all channels. The analysis sequence was as follows: (1) Thresholding according to the Bassoon signal, thereby obtain the Bassoon ROIs (synapses), (2) Erosion of the signals above threshold, to remove small objects that represent noise events, (3) Dilation of the remaining ROIs (true synapses), to return them to synapse size, (4) Analysis of intensity in each channel within the respective ROIs, the average background intensity is subtracted from each value. Background is defined as the signal over the entire image, except for within ROIs. (5) Plotting of intensities per ROI or per image and statistical analysis, (6) obtain Pearson’s correlation coefficients across all pixels in a ROI, for each ROI, between the relevant channels (SNAP tag labeling, Munc13 antibody immunostaining), (7) Plotting of correlation coefficients, statistical analysis. The data in Figure 4 was obtained using ImageJ (version 1.54f). Here, (a) the images were Split to 3 channels (1-DAPI, 2-Synaptophysin, 3-SNAP), (b) filtering and segmentation was applied for channels 1 and 2, (c) a mask image was created and 10 ROIs were randomly selected from each mask and from the background, (d) mean intensity for each ROI was measured within the SNAP channel, and (e) the data was subjected to statistical analysis and plotting.

### Statistics

Statistical analysis was conducted in Prism10. For imaging data, experimental groups were compared against each other using the Mann-Whitney test (Figure 2) or the Kruskal–Wallis test followed by the Dunn’s test for multiple comparisons. For the Electrophysiological data, statistical significance was determined using the non-parametric Mann-Whitney test.

## Data Availability

The data that support the findings of this study are available upon request from the corresponding author.

## ASSOCIATED CONTENT

### SUPPORTING INFORMATION

Supplemental Figures 1-5, Table 1, Generation of an Unc13aSnap knock-in mouse mutant using CRISPR/Cas9 gene editing, Location PCR and genotyping, General Chemistry, SBG-SiR-d12 analysis (PDF)

### Author Contributions

**Maria Kowald**: Conceptualization, Formal analysis, Investigation, Methodology, visualization, Writing – Review and Editing, **Sylvestre Bachollet**: Conceptualization, Formal analysis, Investigation, Methodology, **Fritz Benseler**: Conceptualization, Formal analysis, Investigation, Methodology, Supervision, Writing – Review and Editing, **Maria Steinecker**: Investigation, **Moritz Boll**: Investigation, Formal analysis, **Sofia Kaushik**: Investigation, Methodology, **Tolga Soykan**: Investigation, Methodology, formal analysis, **Sun Siqi**: Formal analysis, Investigation, Writing – Review and Editing, **Ramona Birke**: Investigation, **Dragana Ilic**: Formal analysis, Investigation, Writing – Review and Editing, **Nils Brose**: Conceptualization, Funding acquisition, Supervision, Writing – Review and Editing, **Hanna Hörnberg**: Methodology, Supervision, Writing – Review and Editing, **Martin Lehmann**: Investigation, Methodology, Writing – Review and Editing, **Silvio O. Rizzoli**: Formal analysis, Funding acquisition, visualization, Writing – Review and Editing, **Johannes Broichhagen**: Conceptualization, Formal analysis, Funding acquisition, Investigation, Methodology, Supervision, visualization, Writing – Review and Editing, **Noa Lipstein**: Conceptualization, Formal analysis, Funding acquisition, Investigation, Methodology, Supervision, visualization, Writing – Original draft. All authors have given approval to the final version of the manuscript. / ‡These authors contributed equally.

### Funding Sources

This work was supported by the German Research Foundation Excellence Strategy EXC-2049-390688087 (to N. Lipstein and H. Hörnberg), CRC 1286 “Quantitative Synaptology” project A11 (to N. Lipstein), project A03 (to Silvio O. Rizzoli), and A09 (to N. Brose), the TargetALS foundation (Industry-led collaborative consortia: ‘Correcting Aberrant Splicing of UNC13A as a Therapeutic Approach for ALS and FTD’ to N. Lipstein), and the Einstein Center for Neurosciences (to S. Sun). This project has received funding from the European Union’s Horizon Europe Framework Programme (deuterON, grant agreement no. 101042046 to J. Broichhagen), and from the Chan Zuckerberg Initiative DAF, an advised fund of Silicon Valley Community Foundation, to J. Broichhagen (A Chemical Biology Approach for all-in-one cryoCLEM probes, CC RFA).

### Notes

Conflict of interest: Noa Lipstein is a scientific advisory board member of TRACE Neuroscience Inc. Johannes Broichhagen receives licensing revenue from Celtarys Research, which is unrelated to this project.

## ACKNOWLEDGMENT

The authors would like to thank Christiane Harenberg and Dayana Warnecke from the AGCTLab (MPI for Multidisciplinary Sciences) for expert technical assistance and genotyping. We thank the animal facility of the Max Planck Institute of Multidisciplinary Sciences City Campus for their excellent work towards generating the mouse model, the animal facility of the Max-Delbrück Center Campus Berlin Buch for mouse husbandry, and Blaise Gatin-Fraudet (AG Broichhagen) and Kerstin Steinhagen for technical assistance. We thank the viral core facility, Charite - Universitätsmedizin Berlin for virus production, and Bob Weinberg for the contribution of lentiviral vectors #8454 and #8455 via Addgene.

## ABBREVIATIONS

Munc13-1: protein unc-13 homolog A

SV: synaptic vesicles

BG: *O*^6^-benzylguanine

CA: chloroalkane

ALS: amyotrophic lateral sclerosis

FTD: frontotemporal dementia

SiR: silicon rhodamine

WT: wild type

## REFERENCES

1. Rodriguez, E. A., Campbell, R. E., Lin, J. Y., Lin, M. Z., Miyawaki, A., Palmer, A. E., Shu, X., Zhang, J., Tsien, R. Y. The Growing and Glowing Toolbox of Fluorescent and Photoactive Proteins. Trends Biochem Sci 2017, 42(2), 111-129. DOI: 10.1016/j.tibs.2016.09.010 From NLM Medline.

2. Tsien, R. Y. The green fluorescent protein. Annu Rev Biochem 1998, 67, 509-544. DOI: 10.1146/annurev.biochem.67.1.509 From NLM Medline.

3. Erdmann, R. S., Baguley, S. W., Richens, J. H., Wissner, R. F., Xi, Z., Allgeyer, E. S., Zhong, S., Thompson, A. D., Lowe, N., Butler, R., et al. Labeling Strategies Matter for Super-Resolution Microscopy: A Comparison between HaloTags and SNAP-tags. Cell Chem Biol 2019, 26 (4), 584-592 e586. DOI: 10.1016/j.chembiol.2019.01.003 From NLM Medline.

4. Zhang, D., Chen, Z., Du, Z., Bao, B., Su, N., Chen, X., Ge, Y., Lin, Q., Yang, L., Hua, Y., et al. Design of a palette of SNAP-tag mimics of fluorescent proteins and their use as cell reporters. Cell Discov 2023, 9 (1), 56. DOI: 10.1038/s41421-023-00546-y From NLM PubMed-not-MEDLINE.

5. Keppler, A., Gendreizig, S., Gronemeyer, T., Pick, H., Vogel, H., Johnsson, K. A general method for the covalent labeling of fusion proteins with small molecules in vivo. Nat Biotechnol 2003, 21 (1), 86-89. DOI: 10.1038/nbt765 From NLM Medline.

6. Sun, X., Zhang, A., Baker, B., Sun, L., Howard, A., Buswell, J., Maurel, D., Masharina, A., Johnsson, K., Noren, C. J., et al. Development of SNAP-tag fluorogenic probes for wash-free fluorescence imaging. Chembiochem 2011, 12 (14), 2217-2226. DOI: 10.1002/cbic.201100173 From NLM Medline.

7. Los, G. V., Encell, L. P., McDougall, M. G., Hartzell, D. D., Karassina, N., Zimprich, C., Wood, M. G., Learish, R., Ohana, R. F., Urh, M., et al. HaloTag: a novel protein labeling technology for cell imaging and protein analysis. ACS Chem Biol 2008, 3 (6), 373-382. DOI: 10.1021/cb800025k From NLM Medline.

8. Gautier, A., Juillerat, A., Heinis, C., Correa, I. R., Jr., Kindermann, M., Beaufils, F., Johnsson, K. An engineered protein tag for multiprotein labeling in living cells. Chem Biol 2008, 15 (2), 128-136. DOI: 10.1016/j.chembiol.2008.01.007 From NLM Medline.

9. Masch, J. M., Steffens, H., Fischer, J., Engelhardt, J., Hubrich, J., Keller-Findeisen, J., D’Este, E., Urban, N. T., Grant, S. G. N., Sahl, S. J., et al. Robust nanoscopy of a synaptic protein in living mice by organic-fluorophore labeling. Proc Natl Acad Sci U S A 2018, 115 (34), E8047-E8056. DOI: 10.1073/pnas.1807104115 From NLM Medline.

10. Sahl, S. J., Hell, S. W., Jakobs, S. Fluorescence nanoscopy in cell biology. Nat Rev Mol Cell Biol 2017, 18(11), 685-701. DOI: 10.1038/nrm.2017.71 From NLM Medline.

11. Ries, J., Kaplan, C., Platonova, E., Eghlidi, H., Ewers, H. A simple, versatile method for GFP-based super-resolution microscopy via nanobodies. Nat Methods 2012, 9 (6), 582-584. DOI: 10.1038/nmeth.1991.

12. Bodor, D. L., Rodriguez, M. G., Moreno, N., Jansen, L. E. Analysis of protein turnover by quantitative SNAP-based pulse-chase imaging. Curr Protoc Cell Biol 2012, *Chapter 8*, Unit8 8. DOI: 10.1002/0471143030.cb0808s55 From NLM Medline.

13. Poc, P., Gutzeit, V. A., Ast, J., Lee, J., Jones, B. J., D’Este, E., Mathes, B., Lehmann, M., Hodson, D. J., Levitz, J., Broichhagen, J. Interrogating surface versus intracellular transmembrane receptor populations using cell-impermeable SNAP-tag substrates. Chem Sci 2020, 11(30), 7871-7883. DOI: 10.1039/d0sc02794d From NLM PubMed-not-MEDLINE.

14. Neukam, M., Sala, P., Brunner, A. D., Ganss, K., Palladini, A., Grzybek, M., Topcheva, O., Vasiljevic, J., Broichhagen, J., Johnsson, K., et al. Purification of time-resolved insulin granules reveals proteomic and lipidomic changes during granule aging. Cell Rep 2024, 43 (3), 113836. DOI: 10.1016/j.celrep.2024.113836 From NLM Medline.

15. Yim, W. W., Yamamoto, H., Mizushima, N. A pulse-chasable reporter processing assay for mammalian autophagic flux with HaloTag. Elife 2022, 11, e78923. DOI: 10.7554/eLife.78923 From NLM Medline.

16. Frei, M. S., Tarnawski, M., Roberti, M. J., Koch, B., Hiblot, J., Johnsson, K. Engineered HaloTag variants for fluorescence lifetime multiplexing. Nat Methods 2022, 19 (1), 65-70. DOI: 10.1038/s41592-021-01341-x From NLM Medline.

17. Kühn, S., Nasufovic, V., Wilhelm, J., Kompa, J., de Lange, E. M. F., Lin, Y.-H., Egoldt, C., Fischer, J., Lennoi, A., Tarnawski, M., et al. SNAP-tag2: faster and brighter protein labeling. bioRxiv 2024, 2024.2008.2028.610127. DOI: 10.1101/2024.08.28.610127.

18. Brose, N., Hofmann, K., Hata, Y., Sudhof, T. C. Mammalian homologues of Caenorhabditis elegans unc-13 gene define novel family of C2-domain proteins. J Biol Chem 1995, 270 (42), 25273-25280.

19. Dittman, J. S. Unc13: a multifunctional synaptic marvel. Curr Opin Neurobiol 2019, 57, 17-25. DOI: 10.1016/j.conb.2018.12.011.

20. Augustin, I., Betz, A., Herrmann, C., Jo, T., Brose, N. Differential expression of two novel Munc13 proteins in rat brain. Biochem J 1999, 337 (Pt 3), 363-371.

21. Man, K. N., Imig, C., Walter, A. M., Pinheiro, P. S., Stevens, D. R., Rettig, J., Sorensen, J. B., Cooper, B. H., Brose, N., Wojcik, S. M. Identification of a Munc13-sensitive step in chromaffin cell large dense-core vesicle exocytosis. Elife 2015, 4. DOI: 10.7554/eLife.10635.

22. Brunger, A. T., Choi, U. B., Lai, Y., Leitz, J., Zhou, Q. Molecular Mechanisms of Fast Neurotransmitter Release. Annu Rev Biophys 2018, 47, 469-497. DOI: 10.1146/annurev-biophys-070816-034117.

23. Imig, C., Min, S. W., Krinner, S., Arancillo, M., Rosenmund, C., Sudhof, T. C., Rhee, J., Brose, N., Cooper, B. H. The morphological and molecular nature of synaptic vesicle priming at presynaptic active zones. Neuron 2014, 84 (2), 416-431. DOI: 10.1016/j.neuron.2014.10.009.

24. Lipstein, N., Chang, S., Lin, K. H., Lopez-Murcia, F. J., Neher, E., Taschenberger, H., Brose, N. Munc13-1 is a Ca(2+)-phospholipid-dependent vesicle priming hub that shapes synaptic short-term plasticity and enables sustained neurotransmission. Neuron 2021, 109 (24), 3980-4000 e3987. DOI: 10.1016/j.neuron.2021.09.054 From NLM Medline.

25. Lipstein, N., Sakaba, T., Cooper, B. H., Lin, K. H., Strenzke, N., Ashery, U., Rhee, J. S., Taschenberger, H., Neher, E., Brose, N. Dynamic control of synaptic vesicle replenishment and short-term plasticity by Ca(2+)-calmodulin-Munc13-1 signaling. Neuron 2013, 79 (1), 82-96. DOI: 10.1016/j.neuron.2013.05.011.

26. Augustin, I., Rosenmund, C., Sudhof, T. C., Brose, N. Munc13-1 is essential for fusion competence of glutamatergic synaptic vesicles. Nature 1999, 400 (6743), 457-461. DOI: 10.1038/22768.

27. Basu, J., Betz, A., Brose, N., Rosenmund, C. Munc13-1 C1 domain activation lowers the energy barrier for synaptic vesicle fusion. J Neurosci 2007, 27 (5), 1200-1210. DOI: 10.1523/JNEUROSCI.4908-06.2007.

28. Junge, H. J., Rhee, J. S., Jahn, O., Varoqueaux, F., Spiess, J., Waxham, M. N., Rosenmund, C., Brose, N. Calmodulin and Munc13 form a Ca2+ sensor/effector complex that controls short-term synaptic plasticity. Cell 2004, 118 (3), 389-401. DOI: 10.1016/j.cell.2004.06.029.

29. Rhee, J. S., Betz, A., Pyott, S., Reim, K., Varoqueaux, F., Augustin, I., Hesse, D., Sudhof, T. C., Takahashi, M., Rosenmund, C., Brose, N. Beta phorbol ester- and diacylglycerol-induced augmentation of transmitter release is mediated by Munc13s and not by PKCs. Cell 2002, 108 (1), 121-133.

30. (30) Rosenmund, C., Sigler, A., Augustin, I., Reim, K., Brose, N., Rhee, J. S. Differential control of vesicle priming and short-term plasticity by Munc13 isoforms. Neuron 2002, 33 (3), 411-424.

31. Shin, O. H., Lu, J., Rhee, J. S., Tomchick, D. R., Pang, Z. P., Wojcik, S. M., Camacho-Perez, M., Brose, N., Machius, M., Rizo, J., et al. Munc13 C2B domain is an activity-dependent Ca2+ regulator of synaptic exocytosis. Nat Struct Mol Biol 2010, 17 (3), 280-288. DOI: 10.1038/nsmb.1758.

32. Varoqueaux, F., Sigler, A., Rhee, J. S., Brose, N., Enk, C., Reim, K., Rosenmund, C. Total arrest of spontaneous and evoked synaptic transmission but normal synaptogenesis in the absence of Munc13-mediated vesicle priming. Proc Natl Acad Sci U S A 2002, 99 (13), 9037-9042. DOI: 10.1073/pnas.122623799.

33. Lipstein, N., Verhoeven-Duif, N. M., Michelassi, F. E., Calloway, N., van Hasselt, P. M., Pienkowska, K., van Haaften, G., van Haelst, M. M., van Empelen, R., Cuppen, I., et al. Synaptic UNC13A protein variant causes increased neurotransmission and dyskinetic movement disorder. J Clin Invest 2017, 127 (3), 1005-1018. DOI: 10.1172/JCI90259.

34. Akiyama, T., Koike, Y., Petrucelli, L., Gitler, A. D. Cracking the cryptic code in amyotrophic lateral sclerosis and frontotemporal dementia: Towards therapeutic targets and biomarkers. Clin Transl Med 2022, 12 (5), e818. DOI: 10.1002/ctm2.818 From NLM Medline.

35. van Rheenen, W., van der Spek, R. A. A., Bakker, M. K., van Vugt, J., Hop, P. J., Zwamborn, R. A. J., de Klein, N., Westra, H. J., Bakker, O. B., Deelen, P., et al. Common and rare variant association analyses in amyotrophic lateral sclerosis identify 15 risk loci with distinct genetic architectures and neuron-specific biology. Nat Genet 2021, 53 (12), 1636-1648. DOI: 10.1038/s41588-021-00973-1 From NLM Medline.

36. Bohme, M. A., Beis, C., Reddy-Alla, S., Reynolds, E., Mampell, M. M., Grasskamp, A. T., Lutzkendorf, J., Bergeron, D. D., Driller, J. H., Babikir, H., et al. Active zone scaffolds differentially accumulate Unc13 isoforms to tune Ca(2+) channel-vesicle coupling. Nat Neurosci 2016, 19 (10), 1311-1320. DOI: 10.1038/nn.4364.

37. Reddy-Alla, S., Bohme, M. A., Reynolds, E., Beis, C., Grasskamp, A. T., Mampell, M. M., Maglione, M., Jusyte, M., Rey, U., Babikir, H., et al. Stable Positioning of Unc13 Restricts Synaptic Vesicle Fusion to Defined Release Sites to Promote Synchronous Neurotransmission. Neuron 2017, 95 (6), 1350-1364 e1312. DOI: 10.1016/j.neuron.2017.08.016.

38. Bohme, M. A., McCarthy, A. W., Grasskamp, A. T., Beuschel, C. B., Goel, P., Jusyte, M., Laber, D., Huang, S., Rey, U., Petzoldt, A. G., et al. Rapid active zone remodeling consolidates presynaptic potentiation. Nat Commun 2019, 10 (1), 1085. DOI: 10.1038/s41467-019-08977-6.

39. Mrestani, A., Pauli, M., Kollmannsberger, P., Repp, F., Kittel, R. J., Eilers, J., Doose, S., Sauer, M., Siren, A. L., Heckmann, M., Paul, M. M. Active zone compaction correlates with presynaptic homeostatic potentiation. Cell Rep 2021, 37 (1), 109770. DOI: 10.1016/j.celrep.2021.109770 From NLM Medline.

40. Sakamoto, H., Ariyoshi, T., Kimpara, N., Sugao, K., Taiko, I., Takikawa, K., Asanuma, D., Namiki, S., Hirose, K. Synaptic weight set by Munc13-1 supramolecular assemblies. Nat Neurosci 2018, 21 (1), 41-49. DOI: 10.1038/s41593-017-0041-9.

41. Li, F., Grushin, K., Coleman, J., Pincet, F., Rothman, J. E. Diacylglycerol-dependent hexamers of the SNARE-assembling chaperone Munc13-1 cooperatively bind vesicles. Proc Natl Acad Sci U S A 2023, 120 (44), e2306086120. DOI: 10.1073/pnas.2306086120 From NLM Medline.

42. Grushin, K., Kalyana Sundaram, R. V., Sindelar, C. V., Rothman, J. E. Munc13 structural transitions and oligomers that may choreograph successive stages in vesicle priming for neurotransmitter release. Proc Natl Acad Sci U S A 2022, 119 (7), e2121259119. DOI: 10.1073/pnas.2121259119 From NLM Medline.

43. Wilhelm, B. G., Mandad, S., Truckenbrodt, S., Krohnert, K., Schafer, C., Rammner, B., Koo, S. J., Classen, G. A., Krauss, M., Haucke, V., et al. Composition of isolated synaptic boutons reveals the amounts of vesicle trafficking proteins. Science 2014, 344 (6187), 1023-1028. DOI: 10.1126/science.1252884.

44. Grimm, J. B., English, B. P., Chen, J., Slaughter, J. P., Zhang, Z., Revyakin, A., Patel, R., Macklin, J. J., Normanno, D., Singer, R. H., et al. A general method to improve fluorophores for live-cell and single-molecule microscopy. Nat Methods 2015, 12 (3), 244-250, 243 p following 250. DOI: 10.1038/nmeth.3256 From NLM Medline.

45. Rossmann, K., Akkaya, K. C., Poc, P., Charbonnier, C., Eichhorst, J., Gonschior, H., Valavalkar, A., Wendler, N., Cordes, T., Dietzek-Ivansic, B., et al. N-Methyl deuterated rhodamines for protein labelling in sensitive fluorescence microscopy. Chem Sci 2022, 13 (29), 8605-8617. DOI: 10.1039/d1sc06466e From NLM PubMed-not-MEDLINE.

46. Kalla, S., Stern, M., Basu, J., Varoqueaux, F., Reim, K., Rosenmund, C., Ziv, N. E., Brose, N. Molecular dynamics of a presynaptic active zone protein studied in Munc13-1-enhanced yellow fluorescent protein knock-in mutant mice. J Neurosci 2006, 26 (50), 13054-13066. DOI: 10.1523/JNEUROSCI.4330-06.2006.

47. Abramson, J., Adler, J., Dunger, J., Evans, R., Green, T., Pritzel, A., Ronneberger, O., Willmore, L., Ballard, A. J., Bambrick, J., et al. Accurate structure prediction of biomolecular interactions with AlphaFold 3. Nature 2024, 630 (8016), 493-500. DOI: 10.1038/s41586-024-07487-w From NLM Medline.

48. Padmanarayana, M., Liu, H., Michelassi, F., Li, L., Betensky, D., Dominguez, M. J., Sutton, R. B., Hu, Z., Dittman, J. S. A unique C2 domain at the C terminus of Munc13 promotes synaptic vesicle priming. Proc Natl Acad Sci U S A 2021, 118 (11), e2016276118. DOI: 10.1073/pnas.2016276118 From NLM Medline.

49. Stevens, D. R., Wu, Z. X., Matti, U., Junge, H. J., Schirra, C., Becherer, U., Wojcik, S. M., Brose, N., Rettig, J. Identification of the minimal protein domain required for priming activity of Munc13-1. Curr Biol 2005, 15 (24), 2243-2248. DOI: 10.1016/j.cub.2005.10.055.

50. Quade, B., Camacho, M., Zhao, X., Orlando, M., Trimbuch, T., Xu, J., Li, W., Nicastro, D., Rosenmund, C., Rizo, J. Membrane bridging by Munc13-1 is crucial for neurotransmitter release. Elife 2019, 8, e42806. DOI: 10.7554/eLife.42806.

51. Ast, J., Nasteska, D., Fine, N. H. F., Nieves, D. J., Koszegi, Z., Lanoiselee, Y., Cuozzo, F., Viloria, K., Bacon, A., Luu, N. T., et al. Revealing the tissue-level complexity of endogenous glucagon-like peptide-1 receptor expression and signaling. Nat Commun 2023, 14 (1), 301. DOI: 10.1038/s41467-022-35716-1 From NLM Medline.

52. Birke, R., Ast, J., Roosen, D. A., Lee, J., Rossmann, K., Huhn, C., Mathes, B., Lisurek, M., Bushiri, D., Sun, H., et al. Sulfonated red and far-red rhodamines to visualize SNAP- and Halo-tagged cell surface proteins. Org Biomol Chem 2022, 20 (30), 5967-5980. DOI: 10.1039/d1ob02216d From NLM Medline.

53. Dorrbaum, A. R., Alvarez-Castelao, B., Nassim-Assir, B., Langer, J. D., Schuman, E. M. Proteome dynamics during homeostatic scaling in cultured neurons. Elife 2020, 9, e52939. DOI: 10.7554/eLife.52939 From NLM Medline.

54. Cohen, L. D., Zuchman, R., Sorokina, O., Muller, A., Dieterich, D. C., Armstrong, J. D., Ziv, T., Ziv, N. E. Metabolic turnover of synaptic proteins: kinetics, interdependencies and implications for synaptic maintenance. PLoS One 2013, 8 (5), e63191. DOI: 10.1371/journal.pone.0063191 From NLM Medline.

55. Keller, D., Babai, N., Kochubey, O., Han, Y., Markram, H., Schurmann, F., Schneggenburger, R. An Exclusion Zone for Ca2+ Channels around Docked Vesicles Explains Release Control by Multiple Channels at a CNS Synapse. PLoS Comput Biol 2015, 11 (5), e1004253. DOI: 10.1371/journal.pcbi.1004253.

56. Chen, Z., Das, B., Nakamura, Y., DiGregorio, D. A., Young, S. M., Jr. Ca2+ channel to synaptic vesicle distance accounts for the readily releasable pool kinetics at a functionally mature auditory synapse. J Neurosci 2015, 35 (5), 2083-2100. DOI: 10.1523/JNEUROSCI.2753-14.2015.

57. Nakamura, Y., Harada, H., Kamasawa, N., Matsui, K., Rothman, J. S., Shigemoto, R., Silver, R. A., DiGregorio, D. A., Takahashi, T. Nanoscale distribution of presynaptic Ca(2+) channels and its impact on vesicular release during development. Neuron 2015, 85 (1), 145-158. DOI: 10.1016/j.neuron.2014.11.019.

58. Rebola, N., Reva, M., Kirizs, T., Szoboszlay, M., Lorincz, A., Moneron, G., Nusser, Z., DiGregorio, D. A. Distinct Nanoscale Calcium Channel and Synaptic Vesicle Topographies Contribute to the Diversity of Synaptic Function. Neuron 2019, 104 (4), 693-710 e699. DOI: 10.1016/j.neuron.2019.08.014.

59. Bell, M. E., Bourne, J. N., Chirillo, M. A., Mendenhall, J. M., Kuwajima, M., Harris, K. M. Dynamics of nascent and active zone ultrastructure as synapses enlarge during long-term potentiation in mature hippocampus. J Comp Neurol 2014, 522 (17), 3861-3884. DOI: 10.1002/cne.23646 From NLM Medline.

60. Goel, P., Dufour Bergeron, D., Bohme, M. A., Nunnelly, L., Lehmann, M., Buser, C., Walter, A. M., Sigrist, S. J., Dickman, D. Homeostatic scaling of active zone scaffolds maintains global synaptic strength. J Cell Biol 2019, 218 (5), 1706-1724. DOI: 10.1083/jcb.201807165.

61. Fukaya, R., Hirai, H., Sakamoto, H., Hashimotodani, Y., Hirose, K., Sakaba, T. Increased vesicle fusion competence underlies long-term potentiation at hippocampal mossy fiber synapses. Sci Adv 2023, 9 (8), eadd3616. DOI: 10.1126/sciadv.add3616 From NLM Medline.

62. Bulovaite, E., Qiu, Z., Kratschke, M., Zgraj, A., Fricker, D. G., Tuck, E. J., Gokhale, R., Koniaris, B., Jami, S. A., Merino-Serrais, P., et al. A brain atlas of synapse protein lifetime across the mouse lifespan. Neuron 2022, 110 (24), 4057-4073 e4058. DOI: 10.1016/j.neuron.2022.09.009 From NLM Medline.

63. Wegner, W., Mott, A. C., Grant, S. G. N., Steffens, H., Willig, K. I. In vivo STED microscopy visualizes PSD95 sub-structures and morphological changes over several hours in the mouse visual cortex. Sci Rep 2018, 8 (1), 219. DOI: 10.1038/s41598-017-18640-z From NLM Medline.

64. Bosch, P. J., Correa, I. R., Jr., Sonntag, M. H., Ibach, J., Brunsveld, L., Kanger, J. S., Subramaniam, V. Evaluation of fluorophores to label SNAP-tag fused proteins for multicolor single-molecule tracking microscopy in live cells. Biophys J 2014, 107 (4), 803-814. DOI: 10.1016/j.bpj.2014.06.040 From NLM Medline.

65. Kohl, J., Ng, J., Cachero, S., Ciabatti, E., Dolan, M. J., Sutcliffe, B., Tozer, A., Ruehle, S., Krueger, D., Frechter, S., et al. Ultrafast tissue staining with chemical tags. Proc Natl Acad Sci U S A 2014, 111 (36), E3805-3814. DOI: 10.1073/pnas.1411087111 From NLM Medline.

66. Sutcliffe, B., Ng, J., Auer, T. O., Pasche, M., Benton, R., Jefferis, G. S., Cachero, S. Second-Generation Drosophila Chemical Tags: Sensitivity, Versatility, and Speed. Genetics 2017, 205 (4), 1399-1408. DOI: 10.1534/genetics.116.199281 From NLM Medline.

67. Yang, G., de Castro Reis, F., Sundukova, M., Pimpinella, S., Asaro, A., Castaldi, L., Batti, L., Bilbao, D., Reymond, L., Johnsson, K., Heppenstall, P. A. Genetic targeting of chemical indicators in vivo. Nat Methods 2015, 12 (2), 137-139. DOI: 10.1038/nmeth.3207 From NLM Medline.

68. Gutzeit, V. A., Acosta-Ruiz, A., Munguba, H., Hafner, S., Landra-Willm, A., Mathes, B., Mony, J., Yarotski, D., Borjesson, K., Liston, C., et al. A fine-tuned azobenzene for enhanced photopharmacology in vivo. Cell Chem Biol 2021, 28 (11), 1648-1663 e1616. DOI: 10.1016/j.chembiol.2021.02.020 From NLM Medline.

69. Ast, J., Arvaniti, A., Fine, N. H. F., Nasteska, D., Ashford, F. B., Stamataki, Z., Koszegi, Z., Bacon, A., Jones, B. J., Lucey, M. A., et al. Super-resolution microscopy compatible fluorescent probes reveal endogenous glucagon-like peptide-1 receptor distribution and dynamics. Nat Commun 2020, 11 (1), 467. DOI: 10.1038/s41467-020-14309-w From NLM Medline.

70. Shaib, A. H., Chouaib, A. A., Chowdhury, R., Altendorf, J., Mihaylov, D., Zhang, C., Krah, D., Imani, V., Spencer, R. K. W., Georgiev, S. V., et al. One-step nanoscale expansion microscopy reveals individual protein shapes. Nat Biotechnol 2024. DOI: 10.1038/s41587-024-02431-9 From NLM Publisher.

71. Schmidt, R., Weihs, T., Wurm, C. A., Jansen, I., Rehman, J., Sahl, S. J., Hell, S. W. MINFLUX nanometer-scale 3D imaging and microsecond-range tracking on a common fluorescence microscope. Nat Commun 2021, 12 (1), 1478. DOI: 10.1038/s41467-021-21652-z From NLM Medline.

72. Truckenbrodt, S., Maidorn, M., Crzan, D., Wildhagen, H., Kabatas, S., Rizzoli, S. O. X10 expansion microscopy enables 25-nm resolution on conventional microscopes. EMBO Rep 2018, 19 (9), e45836. DOI: 10.15252/embr.201845836.

73. Chen, F., Tillberg, P. W., Boyden, E. S. Optical imaging. Expansion microscopy. Science 2015, 347 (6221), 543-548. DOI: 10.1126/science.1260088 From NLM Medline.

74. Balzarotti, F., Eilers, Y., Gwosch, K. C., Gynna, A. H., Westphal, V., Stefani, F. D., Elf, J., Hell, S. W. Nanometer resolution imaging and tracking of fluorescent molecules with minimal photon fluxes. Science 2017, 355 (6325), 606-612. DOI: 10.1126/science.aak9913 From NLM Medline.

75. Tycko, J., Wainberg, M., Marinov, G. K., Ursu, O., Hess, G. T., Ego, B. K., Aradhana, Li, A., Truong, A., Trevino, A. E., et al. Mitigation of off-target toxicity in CRISPR-Cas9 screens for essential non-coding elements. Nat Commun 2019, 10 (1), 4063. DOI: 10.1038/s41467-019-11955-7 From NLM Medline.

76. Doench, J. G., Fusi, N., Sullender, M., Hegde, M., Vaimberg, E. W., Donovan, K. F., Smith, I., Tothova, Z., Wilen, C., Orchard, R., et al. Optimized sgRNA design to maximize activity and minimize off-target effects of CRISPR-Cas9. Nat Biotechnol 2016, 34 (2), 184-191. DOI: 10.1038/nbt.3437 From NLM Medline.

77. Concordet, J. P., Haeussler, M. CRISPOR: intuitive guide selection for CRISPR/Cas9 genome editing experiments and screens. Nucleic Acids Res 2018, 46 (W1), W242-W245. DOI: 10.1093/nar/gky354 From NLM Medline.

78. Haeussler, M., Schonig, K., Eckert, H., Eschstruth, A., Mianne, J., Renaud, J. B., Schneider-Maunoury, S., Shkumatava, A., Teboul, L., Kent, J., et al. Evaluation of off-target and on-target scoring algorithms and integration into the guide RNA selection tool CRISPOR. Genome Biol 2016, 17 (1), 148. DOI: 10.1186/s13059-016-1012-2 From NLM Medline.

79. Biesemann, C., Gronborg, M., Luquet, E., Wichert, S. P., Bernard, V., Bungers, S. R., Cooper, B., Varoqueaux, F., Li, L., Byrne, J. A., et al. Proteomic screening of glutamatergic mouse brain synaptosomes isolated by fluorescence activated sorting. EMBO J 2014, 33 (2), 157-170. DOI: 10.1002/embj.201386120.

80. Burgalossi, A., Jung, S., Man, K. N., Nair, R., Jockusch, W. J., Wojcik, S. M., Brose, N., Rhee, J. S. Analysis of neurotransmitter release mechanisms by photolysis of caged Ca(2)(+) in an autaptic neuron culture system. Nat Protoc 2012, 7 (7), 1351-1365. DOI: 10.1038/nprot.2012.074.

81. Lois, C., Hong, E. J., Pease, S., Brown, E. J., Baltimore, D. Germline transmission and tissue-specific expression of transgenes delivered by lentiviral vectors. Science 2002, 295 (5556), 868-872. DOI: 10.1126/science.1067081 From NLM Medline.

82. Camacho, M., Quade, B., Trimbuch, T., Xu, J., Sari, L., Rizo, J., Rosenmund, C. Control of neurotransmitter release by two distinct membrane-binding faces of the Munc13-1 C(1)C(2)B region. Elife 2021, 10. DOI: 10.7554/eLife.72030 From NLM Medline.

83. Stewart, S. A., Dykxhoorn, D. M., Palliser, D., Mizuno, H., Yu, E. Y., An, D. S., Sabatini, D. M., Chen, I. S., Hahn, W. C., Sharp, P. A., et al. Lentivirus-delivered stable gene silencing by RNAi in primary cells. RNA 2003, 9 (4), 493-501. DOI: 10.1261/rna.2192803 From NLM Medline.

