## Supplementary information for "Endogenous SNAP-tagging of Munc13-1 for monitoring synapse nanoarchitecture"

#### Endogenous tagging of Munc13-1 with a SNAP tag as a tool for monitoring synapse nanoarchitecture

### CONTENT

Supplemental Figures 1-5
Table 1
Generation of an Unc13a$^{SNAP}$ knock-in mouse mutant using CRISPR/Cas9 gene editing
Location PCR and genotyping
General Chemistry
SBG-SiR-d12 analysis

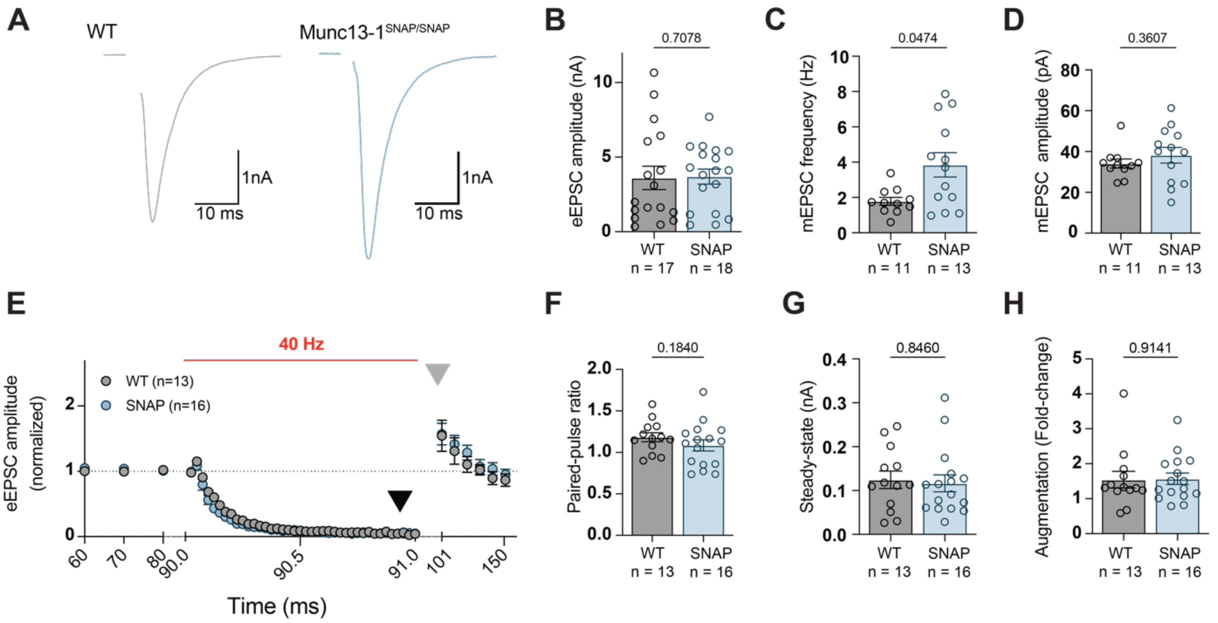


**Figure S1: Electrophysiological analysis of synaptic transmission in Munc13-1$^{SNAP/SNAP}$ neurons. (A)** Example traces of evoked excitatory postsynaptic current (eEPSC) amplitudes recorded in WT (black) and Munc13-1$^{SNAP/SNAP}$ autaptic hippocampal neurons, and (**B**) a plot depicting the averaged initial excitatory eEPSC amplitudes. (**C,D**) Plot depicting the frequency (**C**) and amplitude (**D**) of spontaneous miniature excitatory postsynaptic current (mEPSC) events. (**E**) Averaged and normalized traces of eEPSC amplitudes before, during and after a high frequency action potential train at 40 Hz frequency. (**F**) Paired-pulse ratio (PPR) values, calculated as the ratio of the second to the first eEPSC amplitudes ($eEPSC_2/eEPSC_1$) during the 40 Hz action potential train as in (E). (**G**) Steady-state eEPSC amplitudes calculated as the average of the last three eEPSC amplitudes of the 40 Hz action potential train (see black arrowhead in E), and (**H**) the normalized eEPSC amplitude 10 s after the cessation of the 40 Hz train, reflecting augmentation of the synaptic response following high-frequency activity (see grey arrowhead in E). Circles in **B-D**, **F-H** represent values for individual neurons, respectively. Data in **E** represents the average of the indicated number of individually-recorded neurons. Data was obtained from two independent cultures. Error bars represent mean ± SEM. Statistical analysis was performed by using a Mann-Whitney test.

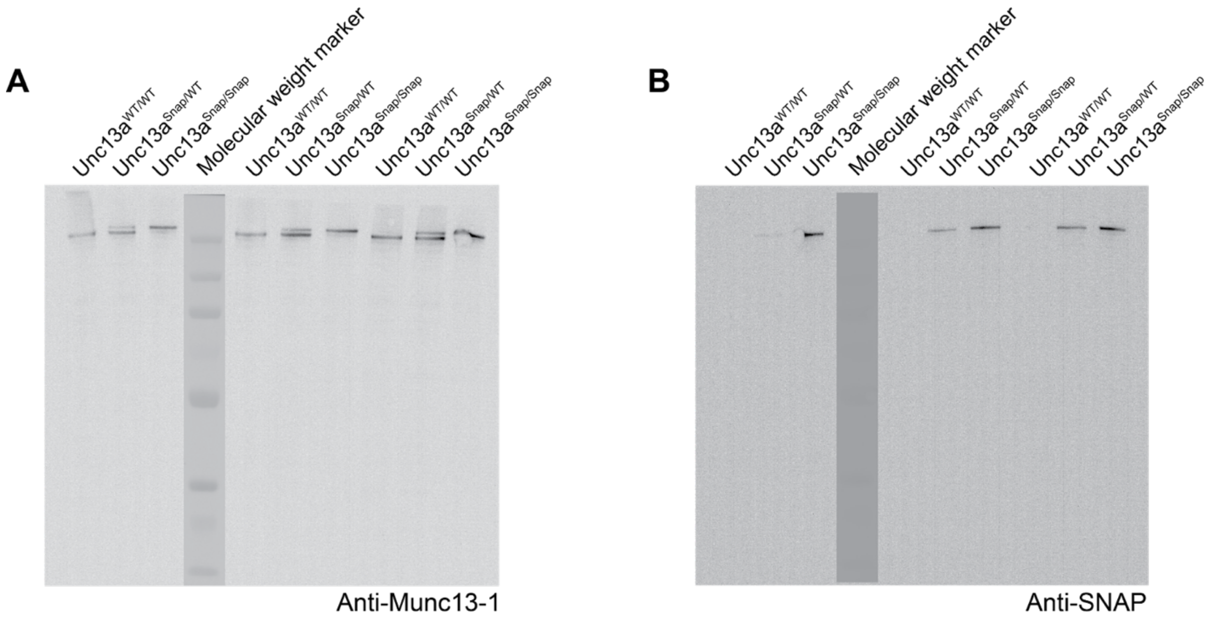


**Figure S2: Endogenous tagging of Munc13-1 with a SNAP tag does not result in truncated protein products.** Uncut Western blot membranes (also used in Figure 1) that were probed with antibodies against **(A)** Munc13-1 or **(B)** the SNAP tag. No truncated protein products are identified in samples from WT, heterozygous and homozygous Unc13a$^{SNAP/SNAP}$ samples.

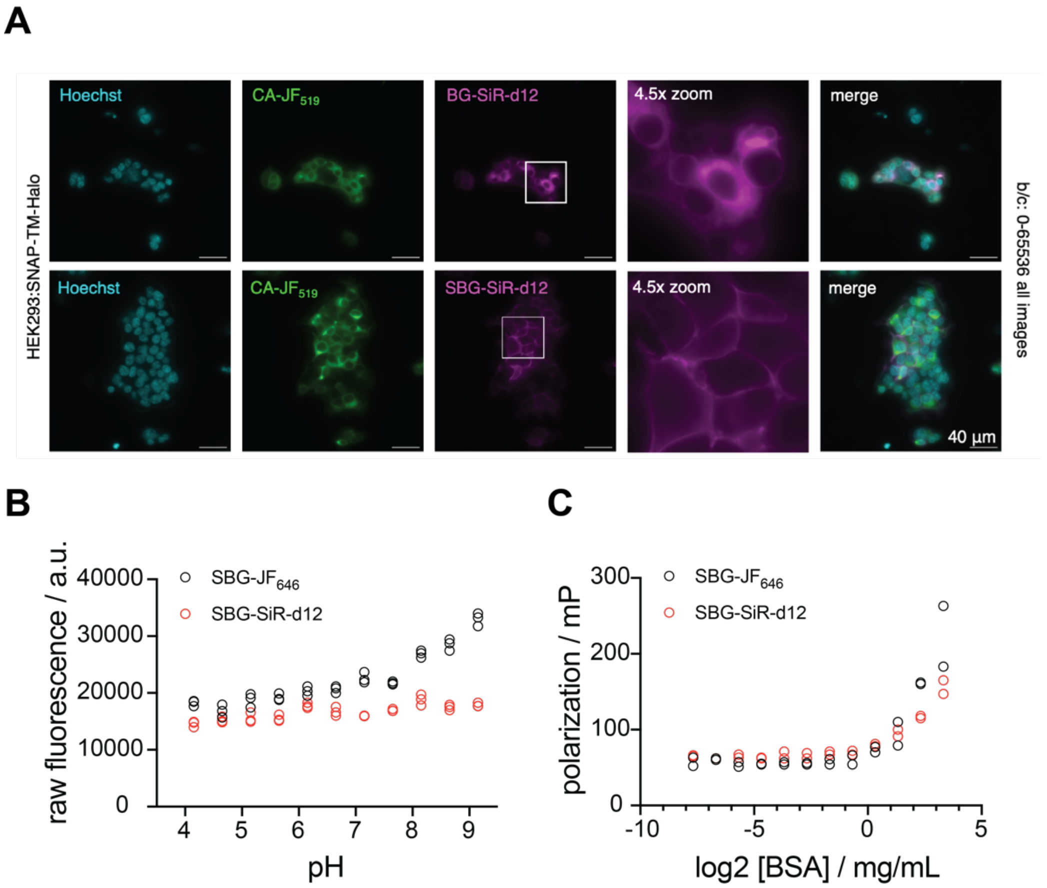


**Figure S3: SBG-SiR-d12 is a membrane-impermeable dye in live cells. (A)** HEK293 cells were transiently transfected with a construct encoding for the expression of SNAP-TM-HTP, a transmembrane construct with an extracellular SNAP tag and an intracellular HaloTag, divided by a single transmembrane helix. Live cells were incubated with CA-$JF_{519}$ targeting the intracellular HaloTag and with either BG-SiR-d12 (upper panel), which can cross membranes (and thus stain intracellular protein pools, for example of newly-translated SNAP-TM-HTP in the endoplasmic reticulum), or with SBG-SiR-d12. Example images of cells with CA-$JF_{519}$ staining clearly demonstrate a diffuse staining by BG-SiR-d12 and a largely membrane-targeted staining by SBG-SiR-d12. White rectangles: indicated areas of zoom-ins. **(B-C)** In vitro measurements of SBG-$JF_{646}$ and SBG-SiR-d12. **(B)** pH sensitivity of SBG-$JF_{646}$ and SBG-SiR-d12 (each 100 nM) shows insensitivity for SBG-SiR-d12 and an increase in fluorescence at more basic conditions for SBG-$JF_{646}$ in phosphate buffer. Measurement performed in triplicates. **(C)** Fluorescence polarization to determine tendency for unspecific binding towards bovine serum albumin (BSA) shows less tendency of SBG-SiR-d12 compared to for SBG-$JF_{646}$. Measurement performed in duplicate.

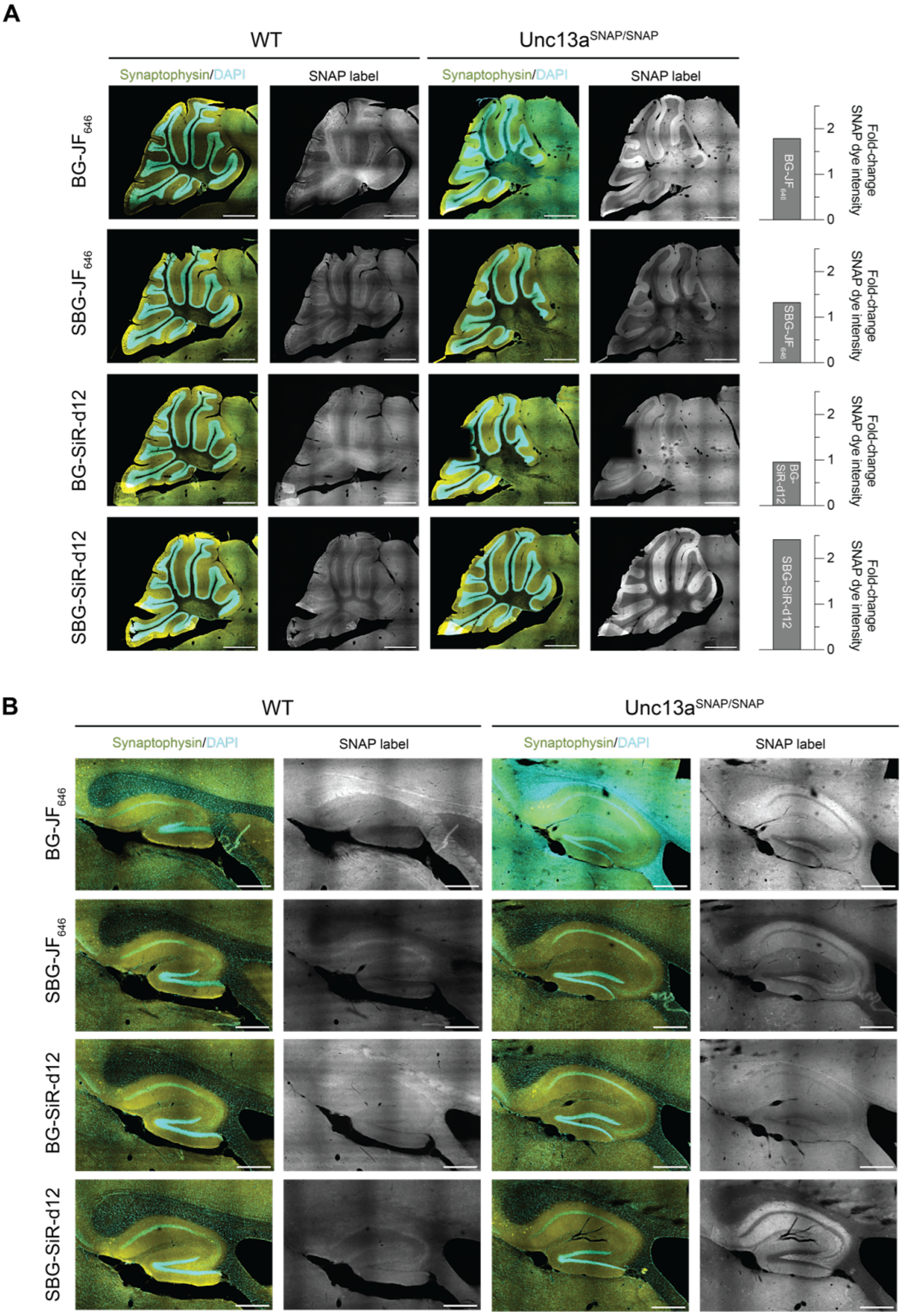


**Figure S4: Labeling of presynaptic terminal in brain sections of Unc13a$^{SNAP/SNAP}$ mice.** Sagittal brain sections from WT and Unc13a$^{SNAP/SNAP}$ mice were stained with either of the following SNAP dyes: BG-JF$_{646}$, SBG-JF$_{646}$, BG-SiR-d12, or SBG- SiR-d12 (as in Figure 4) with an antibody against Synaptophysin 1 (yellow; to stain synapses), and DAPI (cyan), to stain cell nuclei. Example images in the region of the cerebellum (**A**), and the hippocampus (**B**).

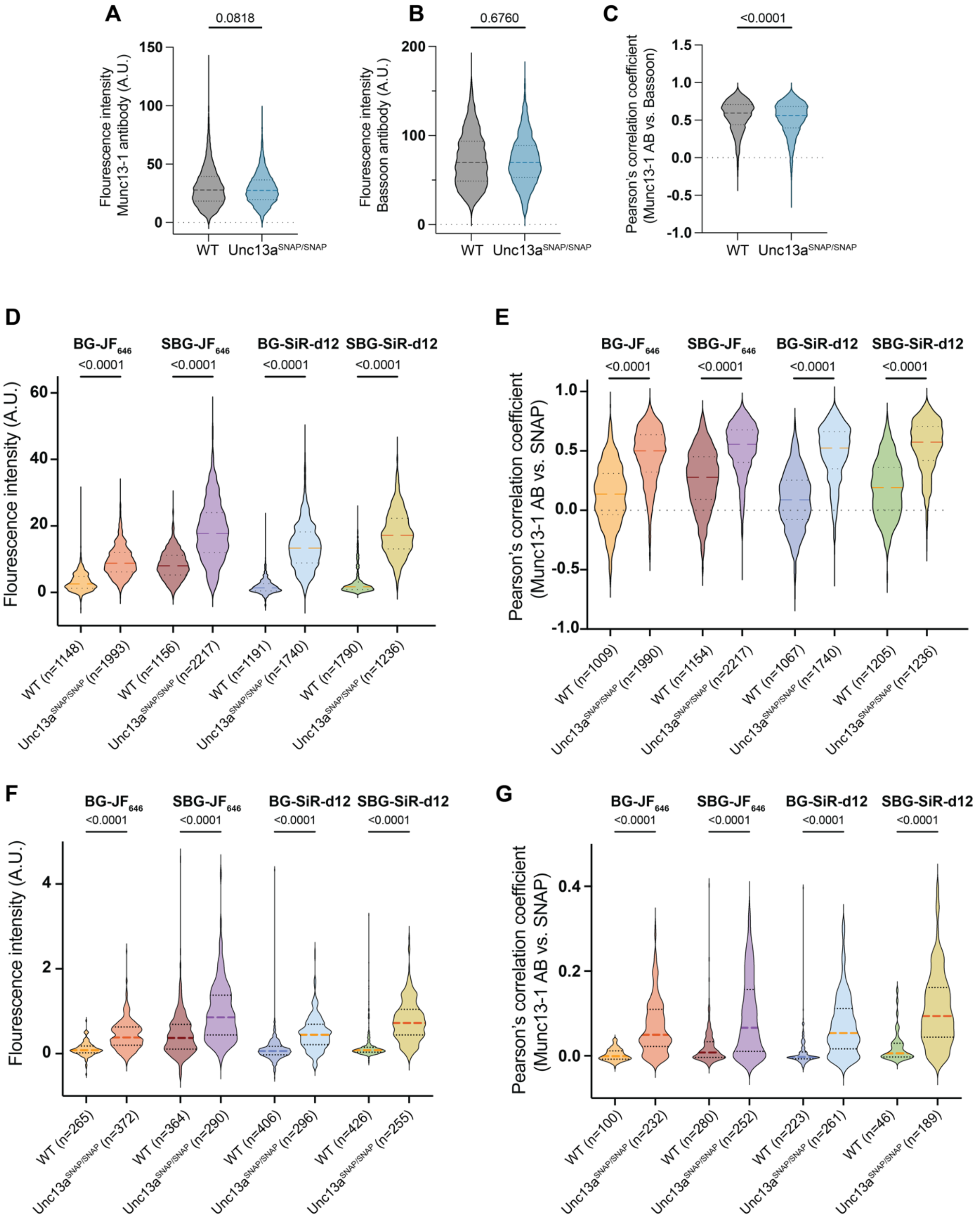


**Figure S5:** Imaging data plotted according to the number of regions of interest (ROIs). **(A-C)** Identical data to the data presented in Figure 2 (C-E), plotted according to the number of ROIs. **(D, E)** Identical data to the data presented in Figure 3, plotted according to the number of ROIs. **(F,G)** Identical data to the data presented in Figure 5 (C,D), plotted by the number of ROIs. In all violin plots, lines represent the median, 25% and 75% quartiles. Statistical significance was evaluated using a two-tailed Mann-Whitney test (A-C) or a two-sided Kruskal-Wallis test followed by Dunn's test for multiple comparisons (D-G). In each panel, data was obtained from three independent experiments.

### *Table 1: Summary of parameters used during microscopy experiments*

| Experiment /Figure | Date | Antigen/Antibody | Dye/ Antibody | Excitation/ Depletion | Laser Power* | Detection |
|---|---|---|---|---|---|---|
| Validation of Munc13-1 localization in hippocampal neurons from Unc13a$^{SNAP}$ mice<br>Leica SP8 STED<br>**Fig. 2**<br>$n_{cultures}$ = 3 per condition<br>$n_{images}$ (WT/Unc13a$^{SNAP/SNAP}$) = 55/54<br>$n_{ROIs}$ (WT/Unc13a$^{SNAP/SNAP}$) = 5285/7186 | 13.6.24<br>17.6.24<br>19.6.24 | Munc13-1/ RRID:AB_887733<br>1:500 | Alexa Fluor™ Plus 594<br>RRID:AB_276 2827<br>1:500 | Confocal: 594 nm | 16%<br>Line Acc. 1 | 604-654 nm |
| | | Bassoon/<br>RRID:AB_2290619<br>1:500 | CF488A<br>RRID:AB_108 53117<br>1:500 | Confocal: 488 nm | 16%<br>Line Acc. 1 | 498-548 nm |
| | | MAP2/<br>RRID:AB_350528<br>1:1000 | Alexa Fluor® 405<br>RRID:AB_289 0171<br>1:500 | Confocal: 405 nm | 100%<br>Line Acc. 1 | 415-465 nm |
| Validation of the SNAP tag functionality in cultured, fixed hippocampal neurons<br>Leica SP8 STED<br>**Fig. 3**<br>$n_{cultures}$ = 3 per condition<br>$n_{images}$ = 10-15 per condition<br>$n_{ROIs}$ = 1000-2200<br>See Figure 3 and S5 for precise n values | 13.6.24<br>17.6.24<br>19.6.24 | Munc13-1/ RRID:AB_887733<br>1:500 | Alexa Fluor™ Plus 594<br>RRID:AB_276 2827<br>1:500 | Confocal: 594 nm | 16%<br>Line Acc. 1 | 604-654 nm |
| | | Bassoon/<br>RRID:AB_2290619<br>1:500 | CF488A<br>RRID:AB_108 53117<br>1:500 | Confocal: 488 nm | 16%<br>Line Acc. 1 | 498-548 nm |
| | | MAP2/<br>RRID:AB_350528<br>1:1000 | Alexa Fluor® 405<br>RRID:AB_289 0171<br>1:500 | Confocal: 405 nm | 100%<br>Line Acc. 1 | 415-465 nm |
| | | BG-SiR-d12/<br>SBG-SiR-d12/<br>BG-JF$_{646}$/<br>SBG-JF$_{646}$ | | Confocal: 647 nm | 80%<br>Line Acc. 1 | 657-707 nm |
| Labeling of presynaptic terminal in brain sections of Unc13a $^{SNAP/SNAP}$ mice at NIKON TiE CSU-X1 Spinning Disc<br>**Fig. 4**<br>$n_{sections}$ (WT/Unc13a$^{SNAP/SNAP}$) = 2 per condition<br>$n_{ROIs}$: 20 per condition | 13.9.24<br>20.1.25 | - | DAPI<br>1:10,000 | 405 nm | 100% (5 mW) | 420-460 nm, 100 ms Exp. |
| | | Synaptophysin 1/<br>RRID:AB_887905<br>1:200 | CF488A<br>RRID:AB_231 3584<br>1:1000 | 488 nm | 100% (7 mW) | 500-550 nm<br>500 ms Exp. |
| | | BG-SiR-d12/<br>SBG-SiR-d12/<br>BG-JF$_{646}$/<br>SBG-JF$_{646}$ | | 640 nm | 100% (12 mW) | 672-744 nm<br>1 s Exp |
| STED microscopy of Munc13-1-SNAP<br>Leica SP8 STED<br>**Fig. 5**<br>$n_{cultures}$ = 3 per condition<br>$n_{images}$ = 6-16 per condition<br>$n_{ROIs}$ = 46-406<br><br>see Figure 5 and S5 for precise n values | 13.6.24<br>17.6.24<br>05.9.24 | Munc13-1/ RRID:AB_887733<br>1:500 | Alexa Fluor™ Plus 594<br>RRID:AB_276 2827<br>1:500 | STED: 594 nm<br>STED Laser: 775 nm | 50%<br>60%<br>Line Acc. 8 | 656-750 nm |
| | | Bassoon/<br>RRID:AB_2290619<br>1:500 | CF488A<br>RRID:AB_108 53117<br>1:500 | STED: 488 nm<br>STED Laser: 592 nm | 40%<br>60%<br>Line Acc. 8 | 498-550 nm |
| | | BG-SiR-d12/<br>SBG-SiR-d12/<br>BG-JF$_{646}$/<br>SBG-JF$_{646}$ | - | STED: 647 nm<br>STED Laser: 775 nm | 100%<br>5%<br>Line Acc. 8 | 656-750 nm |
| Live Imaging of cultured neurons from Unc13a$^{SNAP/SNAP}$<br>Nikon-TiE StedyCON<br>**Fig. 6** | 29.11.24<br>11.12.24<br>17.12.24 | BG-SiR-d12/<br>BG-JF$_{646}$ | - | Confocal: 640 nm<br><br>STED: 640 nm | 100% (51 μW)<br>Line Acc. 1 | 650-700 nm |

| | | | | | | |
|---|---|---|---|---|---|---|
| $n_{cultures}$ = 3 per condition<br>$n_{Images}$ = 60<br>$n_{nanoclusters}$ (Figure 6E)<br>($Unc13a^{SNAP/SNAP}$, BG-$JF_{646}$ vs BG-SiR-d12) = 75/87 | | | | STED Laser 775 nm | 100% (51 µW)<br>Line Acc. 5<br>5% (7 mW) | |
| | | GFP-VGLUT1 | - | Confocal: 488 nm | 100% (13 µW)<br>Line Acc. 1 | 500-550 nm |

* Laserpower for Leica SP8 STED was measured to behave linear, laser power were only measured with a 10 air objective at 100% at 240 uW at 405 nm, 180 uW at 488 nm, 800 uW at 640 nm, 320 mW at 592 nm (CW STED depl.) and 320 mW at 775 nm (pulsed STED depl. 500-550nm; 580-630nm; 650-700nm).

**Generation of an Unc13a$^{Snap}$ knock-in mouse mutant using CRISPR/Cas9 gene editing**

Unc13a *sgRNA1&2 protospacer sequences:*
5'- CTGCGCCCTAGCGCGCGTTT-3' (PAM = CGG)
5'- CCGCCACCGAAACGCGCGCT-3' (PAM = AGG)

Munc13-1_Linker_SNAPtag_HDR1_2335bp:

*HDR1 sequence:*
Unc13a-intronic sequence
Unc13a -Exon44-CDS = BLACK UPPER CASE
Linker = ORANGE UPPER CASE
SNAP-tag = BLUE UPPER CASE
STOP-codon = BOLD UPPER CASE

5'-
ctgctccggccaatagctttgtcatttagagtagttggctcgagtgtggtgcagggtagggatggccgccatacgcggcagaggtg
ggtactggggagatgggatgatagtcgggggctacaggaggctgactggcgtggcccgatcccaggaaactgagcagaagg
cagggtgcagcagggccgggaatggggcgtggccagaatgggtggaaggagtgggcgtggccggagcccaaactcaagct
gctgcgccttaaggatctcagctgcgctgtggggttcacttagtcgcctgccctacgggatgcgctgggcagatcgggtctgggctg
tgggtttcctagtgggcgtggcctagttggagggctttttctgggcgcaacctgggtagcgtggaaggggttctctagccaactttcta
ggagtacgaactactaacgtatgtggcggtgtggtgtgaaccgctcgctggtagctcccctgcgtgacggcgggtgactgggctc
acggtggcctctcgctgagccggtccttgcccacagCTCCCTGAGCGCCGACGCGGCCCCGAGTGCTA
CGAGCTGCAGGTGTGCGTGAAGGACTACTGCTTTGCGCGCGAAGACCGCACGGTGGG
GCTGGCGGTGCTGCAACTGCGAGAGCTGGCCCAGCGCGGGAGCGCCGCGTGCTGGC
TTCCACTCGGCCGCCGCATCCACATGGACGACACGGGGCTCACAGTGCTGCGCATCCT
GTCGCAGCGCAGCAACGATGAGGTGGCCAAGGAATTCGTCAAGCTCAAGTCGGACACG
CGCTCAGCCGAGGAGGGCGGTGCCGCGCCTGCGCCCGGTGGAAGCGGCGGAAGCGG
CGGTAGCATGGACAAAGACTGCGAAATGAAGCGCACCACCCTGGATAGCCCTCTGGGC
AAGCTGGAACTGTCTGGGTGCGAACAGGGCCTGCACCGTATCATCTTCCTGGGCAAAG
GAACATCTGCCGCCGACGCCGTGGAAGTGCCTGCCCCAGCCGCCGTGCTGGGCGGAC
CAGAGCCACTGATGCAGGCCACCGCCTGGCTCAACGCCTACTTTCACCAGCCTGAGGC
CATCGAGGAGTTCCCTGTGCCAGCCCTGCACCACCCAGTGTTCCAGCAGGAGAGCTTT
ACCCGCCAGGTGCTGTGGAAACTGCTGAAAGTGGTGAAGTTCGGAGAGGTCATCAGCT
ACAGCCACCTGGCCGCCCTGGCCGGCAATCCCGCCGCCACCGCCGCCGTGAAAACCG
CCCTGAGCGGAAATCCCGTGCCCATTCTGATCCCCTGCCACCGGGTGGTGCAGGGCG
ACCTGGACGTGGGGGGCTACGAGGGCGGGCTCGCCGTGAAAGAGTGGCTGCTGGCC
CACGAGGGCCACAGACTGGGCAAGCCTGGGCTGGGTTAGcgcgGgtttcgAtggcggggcgggg
cctgggaacggatgagaccccgccttccgtaggccaccgtctctccagaggccacgcccctgcacctcagtggccttggctggt
gaggccctgcctcctgcccctggaaaagccatagagggtccggtgccctgggcatgggctgggcaggcctgcctgggaggcc
gggtcgtagcctctggtgtctgtgcaaaatacagtgtgggatgggctctcaaaagcacacgcgccctcggcgcacacctggggct
gagggcgcctctctccgcggaccccgccctcagcagcgggcccattggcagctggcgacaccctgctcgtgccacacacggtg
ggctgagctccgacgaaaggctcagcccccatcgctgctaaatacccgatttagaatctccctgattcttccccgagggccccac
aggccaggcgccccctccccccaattgaaccggttgccaaacaagtttctgtccttgcccttccttctgggactttagtcggaaccg
gcctagtcaagaagggggggggtgtcccagagccaagggcccccatcccatcttttctccaaatccagtcaggaaataagaga
tcagaaactaggtaggagggagaaagcgagaatatgaacgtgcatttctgagtgtcctgtgcttgtgataaagggatgacccttttt
ctaattggctttgaaccacagcatgtcccctgaatgtcacgtgcagtgacagggtagaattcaggctcttacataggatccaagtttc
atccatggacagggagttcagaggctacattccatacattgagggggggacttgatcccgaccacactggagctcaggagggagt
gatttacactttagggcaacaaacatggctaaaggggaagtggtc**-3'**

**Location PCR and Genotyping**

For genotyping, genomic DNA (gDNA) was isolated from tail biopsies using a genomic DNA isolation kit (Nexttec, #10.924).

For the location PCR, 20 µL reactions were prepared using 1 µL clean gDNA (15-80ng), 4 µL PrimerSet (4 pmol final each), 4 µL 5X Reaction Buffer (Finnzymes #F-524), 0.4 µL PhireHot-Start II Taq DNA Polymerase (Finnzymes #F-122L), 1 µL 10 mM dNTPs (Bioline #DM-515107), 4 µL Hi-Spec Additive (Bioline #HS-014101) and 5.6 µL H2O. Thermocycler parameters: 98°C for 5 min, (98 °C for 45 s, 64 °C for 30 s, 72 °C for 60 s) repeated for 34 cycles, final step at 72 °C for 10 min.

For the diagnostic routine genotyping PCR, 20 µL reactions were prepared using 1 µL clean gDNA (15-80ng), 4 µL PrimerSet (4 pmol final each), 4 µL 5X Reaction Buffer (Biozym #331620XL), 0.2 µL Hot-Start Taq DNA Polymerase (Biozym #331620XL), 1 µL 50 mM MgCl2 (AGCTLab stock) and 9.8 µL H2O. Thermocycler parameters: 96°C for 3 min, (94 °C for 30 s, 62 °C for 60 s, 72 °C for 60 s) repeated for 32 cycles, final step at 72 °C for 7 min.

**Diagnostic, location PCR**

Location1 = 1506bp

5'-AGAAGATGGGCGAGAGGATC-3'
sense_Upstream_HDR1-Munc13-1 (Location 1)

5'-CCCGCCATCGAAACCCGCGCTAACCCAGCCCAGGCTTG-3'
asense-Munc13-1-3’-UTR-SNAP-tag (Location 1)

Location2 = 1564bp

5'-GCGGCGGAAGCGGCGGTAGCATGGACAAAGACTGCGA-3'
sense-Linker-SNAP-tag (Location 2)

5'-ATCTTGGCTCTGTCAGTCAC-3'
asense_Downstream_HDR1-Munc13-1 (Location 2)

**Genotyping strategy**

Primer sequences:

WT band = 162bp

5'-AGCGCAGCAACGATGAGGTG-3'
senseExon44_Munc13-1

5'-GAGAGACGGTGGCCTACGGA-3'
3‘-UTR_Munc13-1

SNAP-tag KI = 225bp

5'-AGCGCAGCAACGATGAGGTG-3'
senseExon44_Munc13-1

5'-CTTTGCCCAGGAAGATGATACGG-3'
asense_SNAP-tag

**General Chemistry**

All chemical reagents and anhydrous solvents for synthesis were purchased from commercial suppliers (Sigma-Aldrich, Fluka, Acros, Fluorochem, TCI) and were used without further purification.

NMR spectra were recorded in deuterated solvents on a Bruker AVANCE III 600 equipped with a CryoProbe calibrated to residual solvent peaks ($^{1}H$ in ppm): MeOD-d4 (3.31). Multiplicities are abbreviated as follows: s = singlet, d = doublet, t = triplet, q = quartet, p = pentet, h = heptet, br = broad, m = multiplet. Coupling constants *J* are reported in Hz. Spectra are reported based on appearance, not on theoretical multiplicities derived from structural information.

UPLC-UV/Vis for purity assessment was performed on an Agilent 1260 Infinity II LC System equipped with Agilent SB- C18 column (1.8 µm, 2.1 × 50 mm). Buffer A: 0.1% FA in $H_2O$ Buffer B: 0.1% FA acetonitrile. The typical gradient was from 10% B for 0.5 min -> gradient to 95% B over 5 mina -> 95% B for 0.5 min -> gradient to 99% B over 1 min with 0.8 mL/min flow. Chromatograms were imported into Graphpad Prism10 and plotted.

High resolution ESI-MS spectra were recorded on a Waters H-class instrument equipped with a quaternary solvent manager, a Waters sample manager-FTN, a Waters PDA detector and a Waters column manager with an Acquity UPLC protein BEH C18 column (1.7 µm, 2.1 mm x 50 mm). Samples were eluted with a flow rate of 0.3 mL/min. The following gradient was used: A: 0.01 % FA in $H_2O$; B: 0.01 % FA in MeCN. 5 % B: 0-1 min; 5 to 95 % B: 1-7min; 95 % B: 7 to 8.5 min. Mass analysis was conducted with a Waters XEVO G2-XS QTof analyzer.

**SBG-SiR-d12 analysis**

$^{1}H$ NMR spectrum:

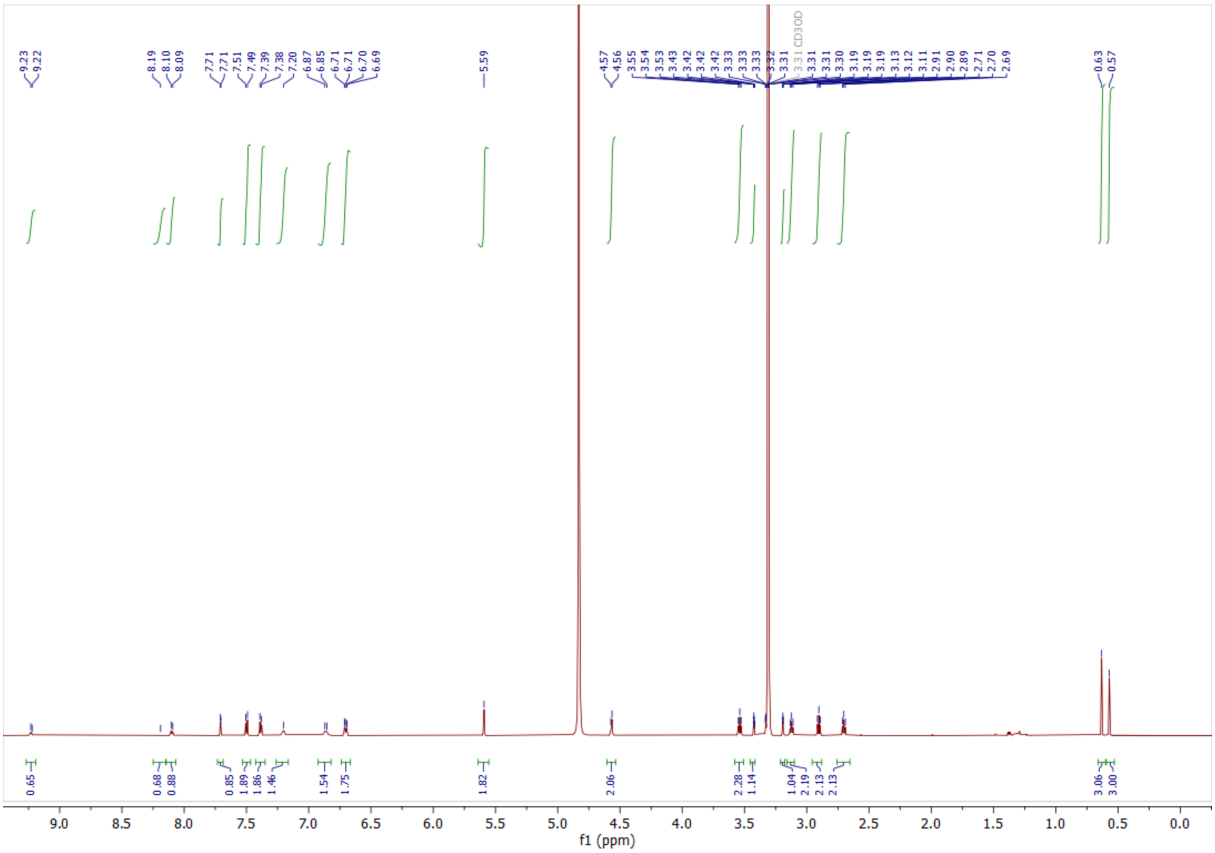
CD3OD
9.0 8.5 8.0 7.5 7.0 6.5 6.0 5.5 5.0 4.5 4.0 3.5 3.0 2.5 2.0 1.5 1.0 0.5 0.0
f1 (ppm)

LCMS trace:

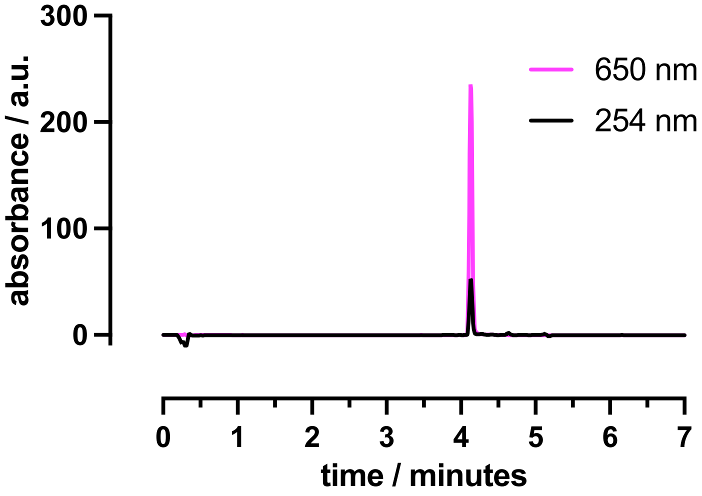
650 nm
254 nm
absorbance / a.u.
300
200
100
0
0 1 2 3 4 5 6 7
time / minutes

Excitation and emission spectra:

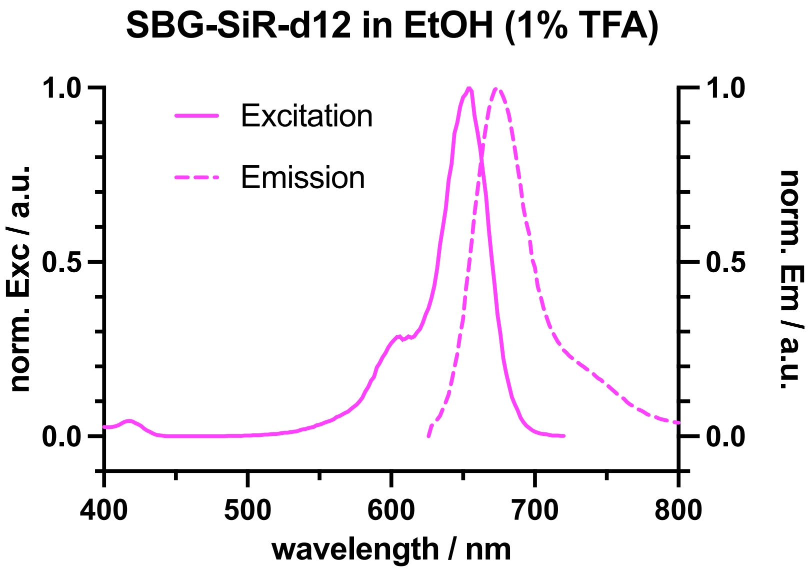


Emission and excitation spectra of SBG_SiR-d12 were determined at a concentration of 100 nM in EtOH + 1% TFA using an infinite 200pro (Tecan) and plotted in Graphpad Prism 10.
